# Direct tensile force activates Adgrl3 in a tethered agonist-dependent manner

**DOI:** 10.64898/2026.02.27.706744

**Authors:** Jesper F. L. Holmkvist, Luis Hamel, Younes F. Barooji, Yin Kwan Chung, Rajesh Regmi, Phillip C. Vejre, Júlia Rosell-Teixidó, Karen L. Martinez, Mette M. Rosenkilde, Poul Martin Bendix, Jonathan A. Javitch, Signe Mathiasen

## Abstract

Adhesion G protein-coupled receptors are proposed to function as mechanosensors, yet whether controlled mechanical force can directly activate receptor signaling in living cells remains unclear. Using optical tweezers, we demonstrate that direct tensile force applied to the N-terminus of the adhesion GPCR Adgrl3 is sufficient to induce G protein recruitment in living cells. Activation is direction-specific, requires a functional tethered agonist, and aligns with coexisting force-driven GAIN-domain conformational changes and dissociation.

---

The adhesion G protein-coupled receptor (GPCR) Adgrl3 participates in transsynaptic interactions, where tensile forces are thought to act across the synaptic cleft, for example during development and synaptic remodeling^1,2^. Indeed, in other cellular systems, adhesion GPCR signaling has been linked to both adhesion and migration contexts where mechanical forces are expected to act^3-7^, yet how such forces are directly coupled to receptor activation in living cells remains unresolved.

Like most adhesion GPCRs, Adgrl3 undergoes autoproteolytic cleavage within the GPCR autoproteolysis-inducing (GAIN) domain, generating a non-covalently associated N-terminal fragment (NTF) and an intramolecular tethered agonist (TA) that remains sequestered within the GAIN domain (Supplementary Fig. 1a); upon exposure, the TA engages the transmembrane core to activate intracellular G protein signaling^8,9^. Single-molecule studies of purified GAIN domains suggest that mechanical force can expose the TA through partial domain unfolding or NTF dissociation^10-12^, but whether these mechanisms operate when the GAIN domain is linked to an intact transmembrane receptor in the fluid plasma membrane of the cell remains unknown. Here, we use optical tweezers to apply controlled mechanical force directly to the extracellular GAIN domain of the adhesion GPCR Adgrl3 while monitoring receptor-proximal G protein recruitment in living cells. We show that direct tensile force is sufficient to activate Adgrl3 in a direction-specific and TA-dependent manner.

**Figure 1.**
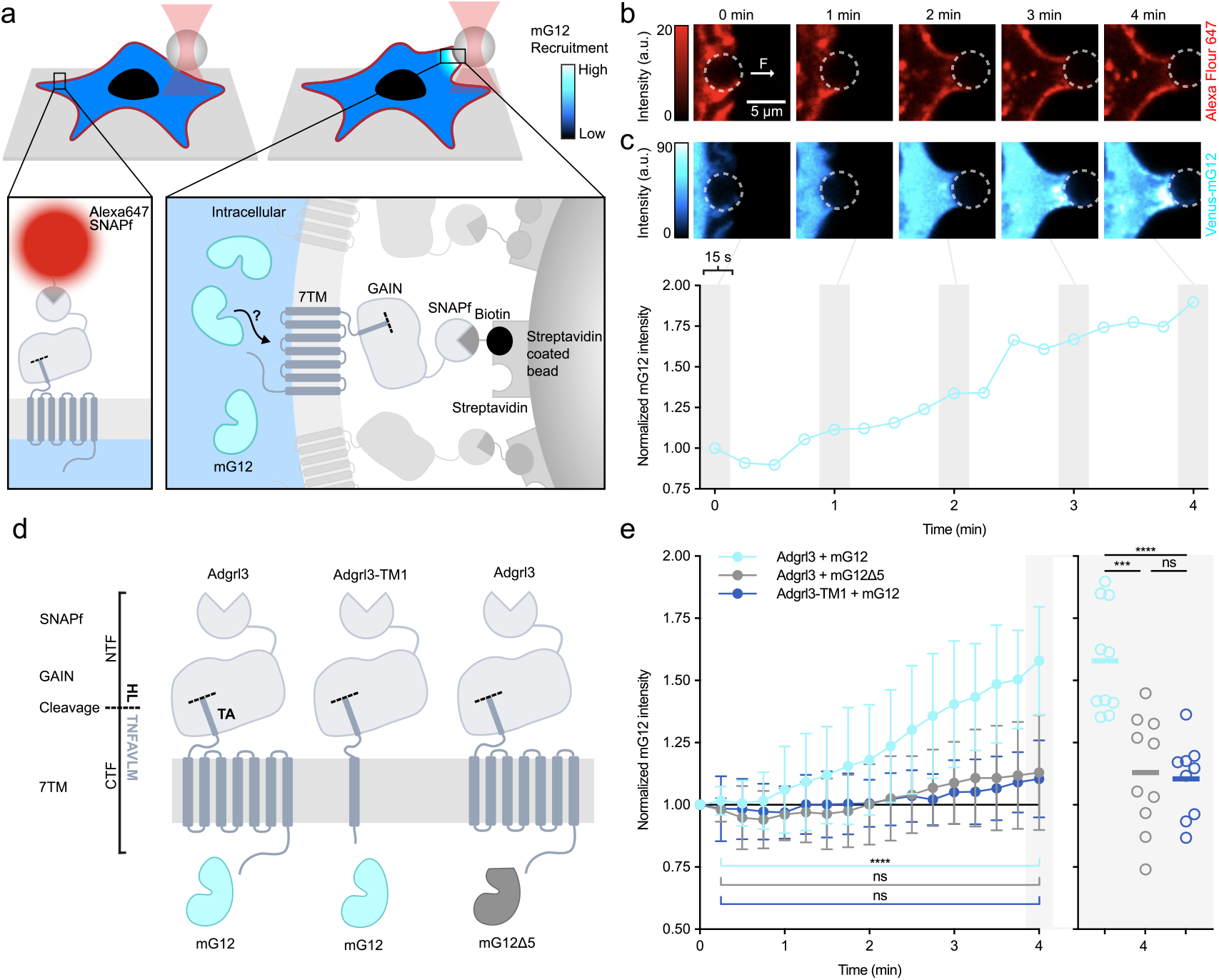
Optical tweezers reveal tensile force-induced activation of Adgrl3. (**a**) Schematic of the experimental setup. HEK293 cells expressing SNAPf-tagged Adgrl3 and Venus-mG12 (mG12) were labeled with BG-Alexa Fluor 647 (AF-647) or BG-biotin. Streptavidin-coated polystyrene beads bound to Adgrl3 were manipulated by optical tweezers. (**b-c**) Confocal micrographs showing AF-647-labeled Adgrl3 (b, red) and mG12 (c, blue) during tensile force application. Circles indicate bead position. (**d**) Schematic of Adgrl3 control constructs used in (e). 7TM, seven transmembrane domain. The dotted line indicates the cleavage site. (**e**) Tensile force induces mG12 recruitment to Adgrl3, but not to Adgrl3-TM1 (which lacks the TM core) or with coupling-deficient mG12Ll5. Data are normalized to t = 0 and shown as mean ± SD (filled symbols) with individual cells at t = 4 min (open symbols); n = 9-10 cells, N = 7-10. Statistics: Unpaired two-tailed t-tests between start and end of force application: ****P < 0.0001, not significant (ns) (Adgrl3 + mG12Ll5, P = 0.0599, Adgrl3-TM1 + mG12, P = 0.0942), one-way ANOVA with Tukey’s multiple comparisons between groups at 4 min: ****P < 0.0001, ***P = 0.0001, ns P = 0.9596.

## Results

To apply controlled mechanical force directly to Adgrl3 in living cells, we engineered an optical tweezer-based force-application assay (Fig. 1a). To isolate force transmission through the GAIN domain, we replaced the N-terminal adhesive domains with a SNAPf-tag, retaining the GAIN domain linked to the transmembrane core (Supplementary Fig. 1a). Full-length Adgrl3 robustly activates the G12/13 signaling pathway^13^, and this construct retained comparable signaling capacity (Supplementary Fig. 1b). We established a monoclonal HEK293 cell line stably expressing this construct (hereafter Adgrl3) and labeled surface-expressed receptors with a 1:1 mixture of BG-Alexa Fluor 647 (AF-647) and BG-biotin to enable both visualization and bead attachment, respectively (Fig. 1a). We then used the optical tweezer to manipulate the bead and apply force directly to Adgrl3, while simultaneously imaging AF-647-labelled receptors by confocal microscopy (Fig. 1b).

We first verified that this experimental design produces specific force load on Adgrl3. At the bead-cell interface, AF-647 fluorescence decreased locally, consistent with displacement of AF-647-labelled receptors by unlabeled, BG-biotin-bound receptors enriched through streptavidin capture (Fig. 1b)^14^. Tensile force was applied by pulling the bead away from the cell, perpendicular to the plasma membrane at a constant speed of 0.05 µm/s. This force application resulted in pronounced cell deformations (Fig. 1b; Supplementary Fig. 2) achieved with an average loading rate of 6 ± 3 pN/s and peak forces of 991 ± 169 pN (Supplementary Fig. 3), consistent with force transmission through multiple receptors at the bead-cell interface^15^. Without BG-biotin, beads failed to engage Adgrl3, producing only thin membrane tethers through nonspecific interactions (Supplementary Fig. 4)^16^, confirming the specificity of receptor-mediated force application.

**Figure 2.**
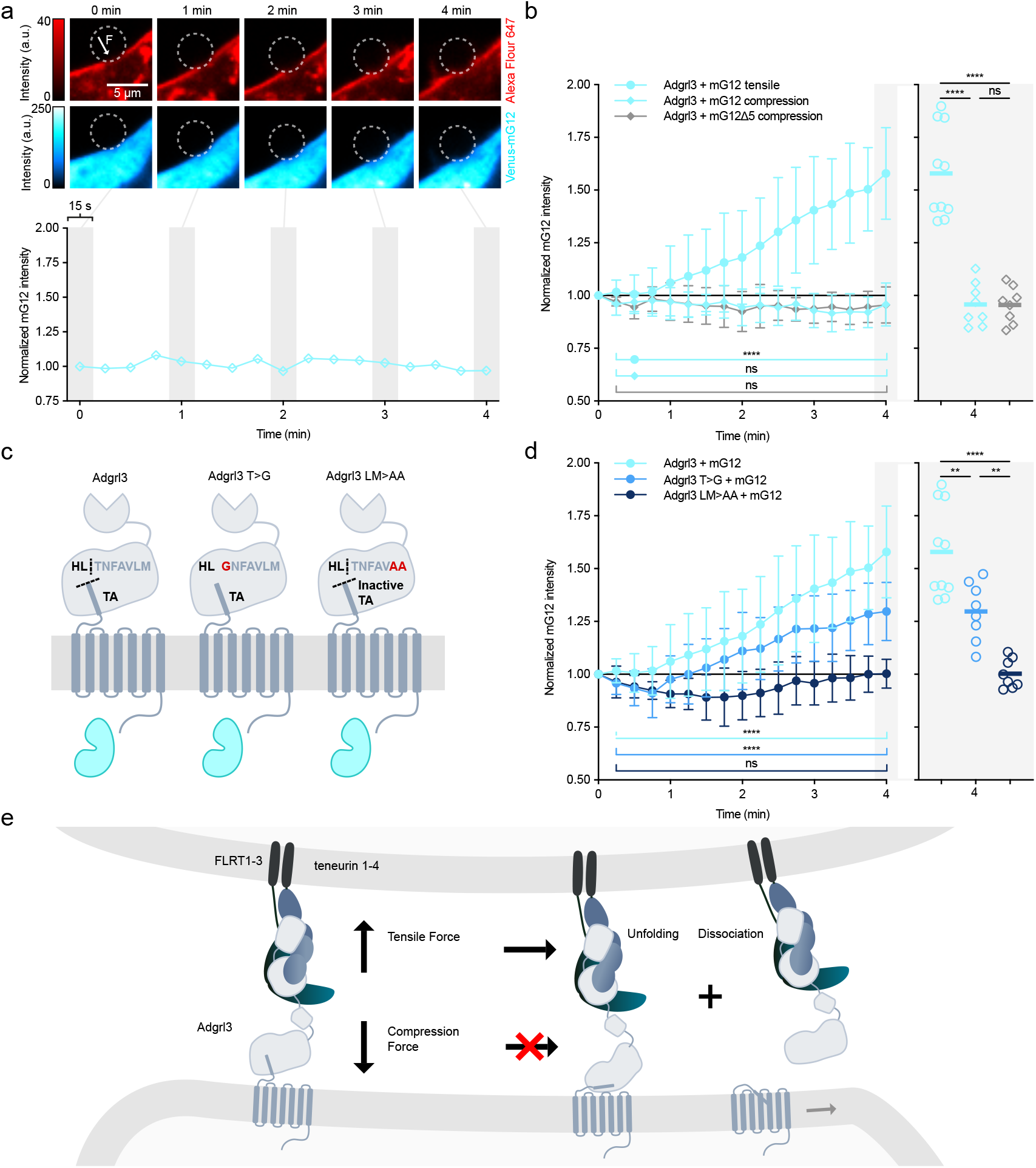
Force direction and TA integrity determine Adgrl3 activation. (**a**) Confocal micrographs showing cell deformation under compressive force application. (**b**) Compressive force does not induce mG12 recruitment. (**c**) Schematic of Adgrl3 TA and cleavage-deficient mutants used in (d): TA-deficient Adgrl3 LM>AA (L928A/M929A) and cleavage-deficient Adgrl3 T>G (T923G). (**d**) Tensile force-induced mG12 recruitment is abolished in Adgrl3 LM>AA and reduced in Adgrl3 T>G. (**e**) Model illustrating tensile force-driven Adgrl3 activation through partial GAIN unfolding and/or N-terminal dissociation. Data in (b) and (d) are normalized to t = 0 and shown as mean ± SD (filled symbols) with individual cells at t = 4 min (open symbols); n = 8-10 cells, N = 6-8. Adgrl3 + mG12 from Fig. 1e is replotted for comparison. Statistics: Unpaired two-tailed t-tests between start and end of force application: (b) ****P < 0.0001, not significant (ns) (Adgrl3 + mG12 compression, P = 0.9747, Adgrl3 + mG12Ll5 compression, P = 0.6028). (d) **** (Adgrl3+mG12, P < 0.0001, Adgrl3 T>G + mG12, P < 0.0001), ns P = 0.3089. One-way ANOVA with Tukey’s multiple comparisons between groups at 4 min: (b) ****(Adgrl3 + mG12 tensile vs. Adgrl3 + mG12Ll5 compression, P < 0.0001, Adgrl3 + mG12 tensile vs. Adgrl3 + mG12 compression, P < 0.0001), ns P = 0.9993. (d) ****P < 0.0001, ** (Adgrl3 + mG12 vs Adgrl3 T>G + mG12, P = 0.0032, Adgrl3 T>G + mG12 vs Adgrl3 LM>AA + mG12, P = 0.0035).

To monitor Adgrl3 activation, we used a minimal G protein alpha subunit (miniG) engineered to remain cytosolic and translocate to the plasma membrane upon GPCR activation^17^. We tested both Venus-miniG12 (mG12) and Venus-miniG13 (mG13) using the endothelin type A receptor (ETA), which activates both G12 and G13, and found that only mG12 robustly responded to receptor activation (Supplementary Fig. 5a). We therefore co-expressed mG12 in cells stably expressing Adgrl3. Upon application of tensile force, mG12 gradually accumulated at the bead-cell interface (Fig. 1c; Supplementary Fig. 2). Quantification of mG12 fluorescence at the interface (Supplementary Fig. 6) revealed a progressive increase over time, reaching a normalized intensity of 1.6 ± 0.2 after 4 min of force application (Fig. 1d-e; Supplementary Fig. 2), demonstrating force-induced recruitment of mG12 by Adgrl3.

To confirm that mG12 recruitment is a consequence of direct mechanical activation of Adgrl3 rather than nonspecific effects, we performed two controls. First, we generated a truncated Adgrl3 lacking the TM core (Adgrl3-TM1; Fig. 1d) and confirmed its absence of signaling activity (Supplementary Fig. 7a). Application of tensile force to cells stably expressing Adgrl3-TM1 at surface levels comparable to Adgrl3 (Supplementary Fig. 8) failed to induce mG12 accumulation (1.10 ± 0.15; Fig. 1e; Supplementary Fig. 9). Second, we used a coupling-deficient miniG construct (mG12Δ5; Fig. 1d), which lacks the C-terminal residues required for productive receptor engagement (Supplementary Fig. 5b). mG12Δ5 was expressed at levels comparable to mG12 (Supplementary Fig. 10). Force application in Adgrl3-expressing cells co-expressing mG12Δ5 did not result in detectable fluorescence accumulation at the bead-cell interface (1.1 ± 0.2; Fig. 1e; Supplementary Fig. 11). These controls demonstrate that tensile force-induced mG12 recruitment is receptor-specific and requires productive Adgrl3-G protein coupling.

While many mechanosensitive receptors respond to tensile forces, compression has also been proposed as a potential activation mechanism for adhesion GPCRs^9^. To test the role of force direction, we subjected cells expressing Adgrl3 and mG12 (or mG12Δ5) to compressive forces by pushing the streptavidin-coated bead toward the cell body (Fig. 2a). Under these conditions, no recruitment was observed with either mG12 (0.96 ± 0.10) or mG12Δ5 (0.96 ± 0.09) (Fig. 2b; Supplementary Fig. 12-13). Although tensile and compressive force application differed in loading rates and peak forces, both converged to comparable plateaus after 4 min. Activation under tensile forces initiates at ∼1 min, corresponding to ∼300 pN, and persists until the plateau (Fig. 1e; Supplementary Fig. 3a). Compression reached similar net forces after ∼1 min and comparable plateau levels without inducing recruitment (Fig. 2b; Supplementary Fig. 3b), indicating that at matched net forces Adgrl3 is not activated by compressive forces.

To test whether mechanical activation of Adgrl3 depends on the TA, we introduced two point-mutations within the TA (Adgrl3 LM>AA; L928A/M929A), previously shown to abolish TA-dependent signaling while preserving receptor cleavage (Fig. 2c, Supplementary Figs. 7b and 14)^18^. Although ensemble measurements implied slightly higher surface expression of Adgrl3 LM>AA, cells selected for force experiments displayed surface expression levels comparable to Adgrl3 (Supplementary Fig. 8). Application of tensile force to cells expressing Adgrl3 LM>AA failed to induce mG12 recruitment after 4 min (1.00 ± 0.07; Fig. 2d; Supplementary Fig. 15), demonstrating that mechanical activation of Adgrl3 requires a functional TA.

Finally, we tested whether receptor cleavage is required for mechanical activation by introducing a cleavage-deficient mutation (Fig. 2c, Adgrl3 T>G; T923G), which retains a functional TA^18^. This mutation abolished cleavage while preserving signaling activity in ensemble assays (Supplementary Figs. 7b and 14). Under tensile force, Adgrl3 T>G showed significantly reduced mG12 recruitment compared to Adgrl3 (1.30 ± 0.14; Fig. 2d; Supplementary Fig. 16), but recruitment was not abolished as it was for Adgrl3 LM>AA. These results indicate that cleavage facilitates, but is not strictly required for, mechanical activation of Adgrl3.

## Discussion

We have developed a new optical tweezer methodology, that enables controlled force application (direction and loading) directly to Adgrl3 at activation-relevant timescales (minutes). Our results support two mechanical activation mechanisms for Adgrl3: an unfolding model, in which the GAIN domain is partly unfolded to expose the TA, and a dissociation model, in which the N-terminal fragment is completely dissociated, leaving the TA fully exposed (Fig. 2e)^5^. These mechanisms are not mutually exclusive. The unfolding model is supported by the finding that cleavage-deficient Adgrl3 can still recruit mG12. The enhanced recruitment observed for cleavage-capable Adgrl3 indicates that dissociation provides a more efficient route to TA exposure. Critically, tensile forces activate Adgrl3, whereas compressive forces at comparable magnitudes do not, indicating that force direction governs activation.

We applied forces in the hundreds of piconewtons, reflecting force regimes needed for global cell deformations transmitted through many bead-receptor connections rather than the force experienced by a single receptor^15^. Single-molecule studies of purified Adgrl3 GAIN domains show that low-piconewton forces can promote unfolding (∼8 pN) or dissociation (∼10-20 pN)^10-12^. Assuming a dissociation force of 15 pN, we estimate at least 65 engaged receptors at the bead-cell contact area when reaching a maximal force of ∼1 nN (Supplementary Fig. 3c). Notably, the bead-cell contact area remains constant throughout cell deformation. Because dissociation releases the NTF from engaged receptors, we anticipate a dynamic turnover at the contact area, with new receptors diffusing in to replace those that have dissociated - consistent with the stable bead-cell interface observed during force application. Additionally, we deformed cells slowly at speeds comparable to physiological traction and migration velocities^19^ (see Methods), allowing cellular viscoelastic relaxation to occur, in contrast to single-molecule experiments with GAIN domains immobilized on rigid glass substrates.

*In vivo*, unfolding and dissociation forces depend on the loading rate^20^. Several factors bias our system towards lower effective loading rates on individual receptors, as compared to single-molecule approaches: multiple receptors sharing the force load, dynamic receptor turnover at the bead-cell contact area, slow deformation velocities, and viscoelastic relaxation. These conditions likely promote unfolding and/or dissociation at low per-receptor forces, consistent with the slip-bond behavior reported for the Adgrl3 GAIN domain^11^, and thus reconcile our high net forces with a model in which piconewton-scale physiological forces drive TA exposure under tension. Resolving the interplay and timing of unfolding and dissociation will require future single-molecule live cell experiments capable of directly capturing GAIN domain unfolding and separation.

## Acknowledgements

This work was supported by Independent Research Fund Denmark, 1054-00086B (S.M.), the Novo Nordisk Foundation grants NNF23OC0081901 (S.M.), NNF23OC008432 (K.L.M., M.M.R, P.M.B., S.M) and NNF20OC0061176 (P.M.B.), NIH grant R21MH112156 (J.A.J.), the Hope for Depression Research Foundation (J.A.J.), and the St. Jude Children’s Research Hospital GPCR Collaborative (J.A.J.).

We thank Dr. Nevin Lambert (Augusta University) for sharing miniG plasmids.

## Author contributions

J.F.L.H., J.A.J. and S.M. wrote the manuscript, with contributions from all the authors; J.F.L.H., J.A.J. and S.M. designed the project; J.F.L.H. performed optical tweezer activity experiments with help from L.H., S.M., and Y.F.B.; J.F.L.H., L.H., and Y.F.B. analyzed data output; L.H., J.F.L.H., and Y.F.B. performed optical tweezer force quantification experiments, and L.H. analyzed data output; Y.K.C., R.R., and P.C.V. performed SRE assays; R.R. performed BRET assays; Y.K.C. performed Western Blots; S.M., R.R., J.F.L.H., and J.R.T. performed cloning of receptor constructs; J.F.L.H., L.H., M.M.R., K.M., P.M.B., J.A.J., and S.M. discussed the experimental findings and interpretation of results; J.A.J. and S.M. supervised the project.

## Declaration of generative Al and Al-assisted technologies in the writing process

Claude (Anthropic), ChatGPT (OpenAI), and Copilot (Microsoft) were used for programming and for language editing, including improvements to clarity, grammar, and consistency. All AI-assisted text and code was reviewed and edited by the authors, who take full responsibility for the content of the manuscript.

## Data availability / Code availability statements

Source data and code supporting this study are available from the corresponding authors upon request.

## Methods

### General cell culture

HEK293 and HEK293T (CRL-3216, ATCC) cell lines were maintained in DMEM (Gibco, Cat. No. 392-0415) supplemented with 10% fetal bovine serum (FBS, Sigma-Aldrich Cat. No. F7524) and 1% penicillin-streptomycin (Thermo Fisher Scientific, Cat. No. 15070063 / in-house vendor “Sterilcentralen”) at 37 °C in a 5% CO_2_-humidified incubator. For the Flp-In T-REx 293 cell lines (Thermo Fisher Scientific, Cat. No. R78007), the culture medium contained additionally 15 µg/mL blasticidin (Thermo Fisher Scientific, Cat. No. R21001 or Avantor, Cat. No. APOSBIB4432) and 100 µg/mL zeocin (Invitrogen, Cat. No. R25001). For the stable cell lines expressing Adgrl3, Adgrl3-TM1, Adgrl3 LM>AA, and Adgrl3 T>G, cells were maintained in medium containing 150 µg/mL hygromycin (Thermo Fisher Scientific, Cat. No. 10687010) and 15 µg/mL blasticidin. All HEK293 cell lines were grown to ∼80% confluency in T25 tissue culture flasks before transfection, tetracycline (Thermo Fisher Scientific, Cat. No. A39246) induction and labelling for microscopy. Cell identity was authenticated by the vendor. Cells were routinely tested to ensure absence of mycoplasma.

### Reagents

Live cell imaging buffer was bought (Thermo Fisher Scientific, Cat. No. A14291DJ) or mixed with following composition: 20 mM HEPES at pH 7.4, 140 mM NaCl, 2.5 mM KCl, 1.8 mM CaCl_2_ and 1.0 mM MgCl_2_. 3× Luciferase substrate solution was mixed with following composition: 15 mM dithiothreitol, 0.6 mM coenzyme A, 0.45 mM ATP, 2.625 mg/mL firefly luciferin (Nanolight, Cat. No. 306-1000), 42 mM Tris base, 127 mM Tris-HCl, 75 mM NaCl, 3 mM MgCl_2_, 7.5% (v/v) Triton X-100^21^.

### Plasmids

Full-length Adgrl3 (NM198702, mouse homolog) cDNA was used to generate the various constructs used in this study. For annotation of amino acids, we use numbering from full-length Adgrl3.

Plasmid construction was made either by restriction enzyme digestion followed by ligation or by Gibson assembly using NEBuilder HiFi DNA Assembly Master Mix (NEB, Cat. No. E2621L). All sequences were confirmed with Eurofins Genomics DNA sequencing service.

cDNAs encoding SNAPf-tagged Adgrl3, Adgrl3-TM1, Adgrl3 LM>AA, and Adgrl3 T>G, were inserted into pcDNA5/FRT/TO (Thermo Fisher Scientific, Cat. No. V652020) for stable cell line generation with the following scheme:

1. pcDNA5/FRT/TO Adgrl3: Directly after the endogenous Adgrl3 signal peptide, a cassette containing a FLAG-tag, a SNAPf-tag, and a flexible linker (GSGSAGD) was inserted upstream of the GAIN domain of Adgrl3 (before position A636). The construct was inserted into the pcDNA5/FRT/TO background vector at restriction sites KpnI-HF and NotI-HF using Gibson assembly.
2. pcDNA5/FRT/TO Adgrl3-TM1: The region after Q974 in (1) was deleted using the one step site directed deletion approach^22^.
3. pcDNA5/FRT/TO Adgrl3 LM>AA: Two alanine substitutions L928A/M929A were introduced in (1) by Gibson assembly and restriction sites NotI-HF and KpnI-HF for insertion into the pcDNA5/FRT/TO background vector.
4. pcDNA5/FRT/TO Adgrl3 T>G: A T923G substitution was introduced in (1) using Gibson assembly and restriction sites NotI-HF and EcoRV-HF for insertion into the pcDNA5/FRT/TO background vector.

Three plasmids expressing miniG proteins were used in the project:

1. NES-Venus-mG13 (mG13): The plasmid was a gift from Professor Nevin Lambert (Augusta University). An N-terminal nuclear export sequence (NES) is placed in front of mG13, and the construct was inserted in the pVenus-C1 background vector^17^.
2. NES-Venus-mG12 (mG12): Fragments of mG12 were amplified by polymerase chain reaction (PCR) using a plasmid carrying NES-Nluc-mG12 (a gift from Professor Nevin Lambert), mG12 were then exchanged for mG13 in the NES-Venus-mG13 vector using BglII and EcoRI-HF restriction sites following the previously described plasmid construction guidelines for the miniG protein series^17^.
3. pcDNA3.1 mG12Δ5: NES-Venus-mG12 lacking the last 5 amino acids were PCR-amplified and inserted into pcDNA3.1+ background vector (Thermo fisher Scientific, Cat. No. V79020) using KpnI-HF and NotI-HF restriction sites.

The remaining plasmids (ETA-Rluc8^17^ and SRE reporter plasmid^23^) used in this study are documented elsewhere.

### Generation of stable HEK293 cell lines

Flp-In T-REx 293 Cell Line was co-transfected with 200 ng of pcDNA5/FRT/TO plasmids encoding the various Adgrl3 constructs and 1,800 ng of the pOG44 Flp-In Recombinant vector (Thermo Fisher Scientific, Cat. No. V600520) using Lipofectamine 2000 (Thermo Fisher Scientific, Cat. No. 11668027) according to the manufacturer’s protocol. Cells were selected in medium in the presence of 250 µg/mL hygromycin and 15 µg/mL blasticidin. After two weeks of antibiotic selection, individual cell colonies were isolated using cloning cylinders (Merck, Cat. No. C2059) and gradually expanded in cell population. For each cell line 4-6 monoclonal cell populations were induced overnight by 1 µg/mL tetracycline and tested for receptor expression by SNAPf surface labelling using membrane-impermeant BG-Alexa Fluor 647 (New England Biolabs, Cat. No. S9136S). Fluorescence was observed using a Leica SP5 confocal microscope (Leica Microsystems). Alexa Fluor 647 was excited by a 633 nm laser, and emission was collected at 650-750 nm. One final clone for each Adgrl3 construct was selected for further experiments, based on visual inspection of cell health and good receptor surface labelling.

### Serum responsive element (SRE) ensemble assay

HEK293T cells were seeded in a 12-well culture plate at a density of 0.3 × 10^6^ cells per well. The following day, cells were transiently co-transfected with 600 ng of SRE reporter plasmid that drives expression of firefly luciferase by SRE-sensitive promoter, and varying amounts of receptor plasmid. 2.33 µL of Lipofectamine 2000 was used per 1 µg of plasmid DNA. After 24 hours of transfection, cells were serum-starved for 6 hours. Cells were then washed twice with HBSS and suspended in 600 µL of HBSS. 80 µL of the suspension was transferred to a well of 96-well polystyrene microplates with flat bottom (Greiner Bio-One, Cat. No. 655074) and was lysed with 40 µL of 3× luciferase substrate solution. Lysates were incubated in darkness for 10 minutes and shaken for 2 minutes. Luminescence was measured by EnVision Multilabel Plate Reader (PerkinElmer).

### Bioluminescence Energy transfer ensemble assay

HEK293T cells were seeded in six-well culture plates at a density of 0.9 x 10^6^ cells per well. The following day, cells were transfected with 3.2 µL Lipofectamine 2000 per 1 µg of cDNA. Cells were transfected with ETA-Rluc8, either one of the miniG plasmids and pcDNA3.1 in a 1:20:20.6 ratio for a total of 2.5 µg in each well of a six-well culture plate. The medium was changed 24 hours after transfection, and after approximately ∼48 h, the cells were washed with HBSS, and two wells were combined for each dose response curve, by resuspension in a total of 1.7 mL HBSS and distributed into 96-well polystyrene microplate at 45 µL per well. Cells were incubated for 7 minutes with ET-1 ligand (Tocris, Cat. No. 1160) before addition of coelenterazine H (Final concentration: 5 µM) (NanoLight Technologies, Cat. No. 301) to reach a final well volume of 100 µL. Rluc8 (donor) emission was read at 480/30 nm, and mVenus (acceptor) emission was read at 535/30 nm, using an EnVision Multilabel microplate reader 30 minutes after ligand addition. The BRET signal was calculated as the ratio of light emitted at 535 nm over that emitted at 480 nm. The ligand-induced BRET ratio was obtained by subtracting baseline BRET (buffer) for each condition.

### Western blotting

Stable cell lines expressing Adgrl3, Adgrl3-TM1, Adgrl3 LM>AA or Adgrl3 T>G, treated with tetracycline (0.25 µg/mL) for overnight receptor induction, were lysed in RIPA buffer (Merck, Cat. No. 20-188) and precleared of insoluble debris by centrifugation. Lysates were mixed with 5× Laemmli buffer and vortexed briefly. 30 µL of samples were loaded onto 4-20% Mini-PROTEAN TGX precast protein gel (BioRad, Cat. No. 4561095) until well separated. Proteins were transferred onto nitrocellulose membranes and subsequently blocked by blocking buffer:phosphate-buffered saline (PBS) (1:1) (PBS-T, Licor, Cat. No. 927-70001, 0.1% (v/v) Tween-20) for 1 hour at room temperature. Membranes were incubated in blocking buffer-diluted α-SNAP-tag (1:1000, New England Biolabs, Cat. No. P9310S) or α-tubulin-β (1:2000, Thermo Fisher Scientific, Cat. No. 322600) overnight at 4°C with shaking. Membranes were then washed twice with PBS-T for 5-minute shaking each at room temperature, followed by incubation in PBS-T-diluted IR680RD-conjugated α-Rabbit (1:15000, Licor, Cat. No. 926-68071) or IR800CW-conjugated α-Mouse (1:15000, Licor, Cat. No. 926-32210) for 1 hour at room temperature with shaking. Membranes were washed twice with PBS for 5-minute shaking each at room temperature. Immunoreactivities were detected using Odyssey XF Dual-Mode Imaging System (Li-Cor), with 700 nm and 800 nm for α-Rabbit and α-Mouse channels respectively. Quantifications on the intensities of Adgrl3 FL and Adgrl3 NTF bands were done by ImageJ with black-white colored blot images. The extent of GAIN domain cleavage was calculated as the percentage fraction of NTF in SNAP-tag immunosignals.

### Microscopy sample preparation

Stable cell lines expressing Adgrl3, Adgrl3-TM1, Adgrl3 LM>AA or Adgrl3 T>G were plated in 3.5 mm round glass bottom dishes (Mattek, Cat. No. P35G-1.5-10-C) in cell culture medium and incubated at 37°C in 5% CO_2_ overnight. Before cell plating, the glass dishes were coated with fibronectin (Sigma-Aldrich, Cat. No. F2006-2MG) by incubating with a 10 µg/mL solution in PBS for 40 minutes and then washed three times with excess PBS. Cells at ∼50% confluency were then transfected overnight with 1 µg mG12 or mG12Δ5 per 2 mL medium using 4 µL of FuGENE HD (Promega, Cat. No. E2311) as directed by the manufacturer’s protocol. Next day, cells at ∼70% confluency were induced with 0.25 µg/mL tetracycline overnight. On the day of experiment cells were labelled in cell culture medium containing 500 ng/mL BG-Alexa Fluor 647 and 500 ng/mL BG-biotin (New England Biolabs, Cat. No. S9110S) for 30 minutes at 37°C in 5% CO_2_ and subsequently incubated with fresh cell culture medium for 40 minutes at 37°C in 5% CO_2_. Immediately before imaging, cells were washed three times with excess imaging buffer, and a final volume of 1 mL imaging buffer was added together with 2.5 µL Streptavidin-coated polystyrene beads (diameter 4.36 µm, Spherotech, Cat. No. SVP-40-5).

For mG12 recruitment experiments (Fig. 1-2, Supplementary Figs. 2, 4, 9, 11, 12, 13, 15 and 16) measurements were performed in an open chamber allowing for 1 mL of buffer reservoir.

For the quantitative force measurements (Supplementary Fig. 3), measurements were done ∼48h after cell seeding, following same procedure as described above, but omitting the mG12 transfection step. Additionally, after labelling and adding the beads, the chambers were sealed using a glass coverslip (#1.5H Glass, Ibidi, Cat. No. 10815) and the excess buffer was removed.

### Optical trapping and confocal imaging

Experiments were performed using the commercially available LUMICKS C-Trap optical tweezers system. A near-infrared trapping laser (1064 nm) was tightly focused through a Nikon 60× water-immersion objective (CFI Plan Apo, NA 1.2) to generate a highly stable gradient optical trap inside the experimental chamber.

A high-precision piezoelectric stage enabled accurate positioning of the trap to capture and move beads inside the microscopy chamber. Confocal fluorescence imaging was recorded simultaneously using two excitation lasers at 488 nm (for excitation of Venus-mG12) and 640 nm (for excitation of Alexa Fluor 647), with emission detected in two channels with a 512/25 nm filter, and a 640 LP filter, respectively. Pixel size was set to 250 nm, and the excitation laser powers were 1.5-2% (0.56-0.76 µW) for the 488 nm laser and 1% (0.94 µW) for the 640 nm laser.

For force detection, forward-scattered light from the trapping laser was collected by an oil-immersion condenser and directed onto a position-sensitive detector (PSD). The PSD recorded bead displacements at a 70 kHz sampling rate, allowing precise quantification of force changes during the experiments. For force measurements, in addition to confocal imaging, bright-field imaging of the sample plane was performed in parallel at 60 frames per second.

### Force application procedure

Force application was initiated by moving the optical trap to bring a trapped bead into contact with the cell. After contact was established, the bead was carefully pushed and moved parallel to the cell surface (∼30 s) to enable firm attachment of the bead via receptor-streptavidin connections. After ∼40 s rest at the bead-cell contact area, an automated 0.05 µm/s trap movement was initiated in a direction perpendicular to the cell surface, either away from the cell (tensile) or towards the cell (compression).

For the tensile experiments, the laser power was increased when the bead began to be displaced from the central trapping region by approximately the bead radius. Increasing the laser power re-positioned the bead within the trap center and ensured that the bead stayed in the linear detection region, which equals the bead radius^24^. The laser power was first increased from 15% to 25% and finally to 35% for the remaining time of the experiment.

The trapping laser powers used, 15, 25 and 35%, corresponding to 380, 640 and 900 mW (Supplementary Fig. 3a) as measured at the focal plane. Local heating induced by the optical trap at the highest power level is smaller than 10°C and likely less due to the bead’s proximity to the glass surface during the experiment^25^. The local temperature-increase decays from the heat source with distance^26-28^ further reducing any potential heat effects on the cells. Local heat gradients cause cells to form blebs^29^ and therefore any cells showing signs of blebbing at any timepoint were discarded.

For the compression experiments, the laser power was kept at 15% throughout the whole measurement to minimize the laser light penetrating the cell. Once a stable deformation was created, the automated movement was stopped, and the stage and trap kept stationary until the end of the measurement. When applying compression forces, there is a probability that the cell morphology pushes the bead in the z-direction and therefore out of the trap focus, which consequently lead to inaccurate force measurements (Supplementary Fig. 3d). These cells were also excluded from the analysis.

### Force quantification

For quantitative force measurements, the trap position was held stationary, and the sample was moved using a stage controller to bring the cell into contact with the trapped bead. Once the receptor-streptavidin connections were established as described above, the stage was automatically moved at a constant speed of 0.05 µm/s. For quantitative force measurements, a single bead was optically trapped initially at 15% (380 mW) laser power and calibration were achieved by automated fitting of a Lorentzian function to the power spectral density of the Brownian bead fluctuations (average spring constant 0.27 ± 0.03 pN/nm, N = 19). The base-line force was set to zero corresponding to a position of the bead at the center of the trap.

To calibrate the trap at the higher laser powers used here, the laser power was first increased to 25% (640 mW) and subsequently to 35% (900 mW) for 5 seconds at each power level, allowing us to record an experimental force offset. This offset was later used for correcting the acquired force profile (Supplementary Fig. 3). To validate that forces remain correct after changing the laser power we performed a control experiment in which a known force was exerted on the bead through a fluid drag by oscillating the fluid surrounding the trapped bead. The hydrodynamic force on the bead is given by:

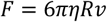

where v is the velocity of the fluid, η is the viscosity of water and R is the radius of the bead. We obtain the correct fluid drag force profile under oscillation at each laser power by subtracting the corresponding measured offset (Supplementary Fig. 3e).

Real-time force data were exported from Bluelake HDF5 files and analyzed using custom scripts in the Pylake (v1.8.0, LUMICKS) Python (version 3.12) package. Force signals were downsampled to 5 Hz for plotting and analysis.

The force offset which takes place when adjusting the laser power to 25% and 35% was respectively subtracted from the measured force (Supplementary Fig. 3). Time t = 0 was defined at the point when the total force was at a minimum, and during the following 4 minutes the peak force and the final relaxation plateau (final 20 s) were quantified. Linear fits using the least square method and an R^2^ > 0.98 criteria were fit to the different force regimes. The mean loading rate applied was calculated within the interval ranging from the force minimum to the end of the stage movement (T), using the following equation:

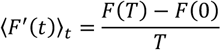

Where *⟨F*^′^(*t*)*⟩*_*t*_ is the average loading rate, and *F*(*0*) and *F*(*T*) are the recorded force acting on the bead at the start and end of the stage movement, respectively.

### Quantification of Venus-mG12 recruitment

Confocal microscopy data were exported from Bluelake HDF5 files using the Pylake Python package and tiff sequences were exported directly from the Bluelake software and analyzed using custom macros and built-in Fiji plugins in the open-source distribution of ImageJ, Fiji (version 2.16.0/1.54p)^30^. First, 4-14 images, depending on the temporal resolution, were averaged to reduce noise, resulting in a temporal resolution of ∼15 s.

To accurately locate the cell outline, the receptor and mG12 intensity channels were merged to include all fluorescence signals, and a mask (ImageJ Default auto-thresholding method (IsoData / iterative intermeans), assuming a dark background) was applied (Supplementary Fig. 6).

The direction of the deformation was specified manually as a straight line, and several parallel lines with a distance of 1 pixel were added, to span the entire width of the bead-cell interface. To track and quantify the mG12 intensity signal at the bead-cell interface at each 15 s timepoint, a 5×5 pixel region was then created at the intersection of the cell outline (mask periphery) and the deformation direction lines. Finally, all pixels in the 5×5 pixel region were combined into one ROI spanning the entire width of the bead-cell interface and ∼3 pixels (750 nm) into the cell. This size captures the bead-membrane interface but excludes global intracellular intensity variations. The pixels outside the mask were excluded, before measuring the average intensity in the mG12 channel of the pixels inside the ROI.

Time t = 0 was defined as the beginning of the deformation for tensile forces and as the start of the automated stage movement for compression forces, and the intensities were normalized to t = 0 using MATLAB (MathWorks, version R2024b).

### Quantification of receptor expression levels across cell lines

To compare receptor surface expression levels across the stable cell lines included in the mG12 recruitment experiments (Fig. 1-2) Fiji was used to draw line scans at ten locations across the cell membrane in the first frame before contact was established with the bead. The intensity profiles were measured (Supplementary Fig. 8a-b) and the mean of the ten maximum intensities from each line scan was used as the expression level for each individual cell.

For ensemble measurements of receptor surface expression (Supplementary Fig. 8f), stable cell lines seeded in a 12-well plate were incubated with or without tetracycline (0.25 µg/mL) overnight for receptor induction. Cells were then incubated with BG-Alexa Fluor 647 for 30 minutes at 37°C and subsequently washed twice with HBSS and suspended in 300 µL of HBSS per 12-well. 50 µL of the suspension was transferred to a well of 96-well black opaque microplate and 3 wells were used per condition. Intensity signals from Alexa Fluor 647 were detected by CLARIOstar plate reader (BMG Labtech) with a built-in protocol exciting at 625-630 nm, and collecting the emission intensity between 630-680 nm, with a dichroic wavelength automatically set as 652.5 nm.

### Quantification of Venus-mG12 expression levels

Three chambers with Adgrl3 LM>AA cells were transfected with mG12 or mG12Δ5. In each chamber ten cells were selected, using the same criteria as for mG12 recruitment experiments: Cells should have both the receptor and mG12 intensity signals, be healthy without any sign of blebbing and have a membrane area available for bead attachment. The ten cells were imaged in the LUMICKS C-Trap under identical confocal imaging conditions, as used in the mG12 recruitment experiments (Fig.1-2), with a 488 nm laser power of 1.75% and 640 nm laser power of 1%. To extract the mean Venus-mG12 intensity per cell, the ten frames were averaged in Fiji, neighboring cells were excluded manually, and all signals outside the cell were then removed using Fiji’s default auto-threshold algorithm (IsoData (Iterative Intermeans) thresholding). Finally, the mean Venus-mG12 intensity per cell was calculated (Supplementary Fig. 10).

### Estimation of deformation velocity

To determine the deformation velocity, we tracked the centroid of the bead-cell surface ROI (see methods ‘Quantification of Venus-mG12 recruitment’ and Supplementary Fig. 6). This allowed us to calculate an average deformation speed:

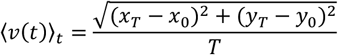

Where (*x*_0_, *y*_0_) and (*x*_*T*_, *y*_*T*_) are the centroid coordinates at the start and end of the deformation movement, respectively. Using this strategy, we arrive at a deformation velocity of 34 ± 7 nm/s (Supplementary Fig. 3f) for the Adgrl3 and mG12 expressing cells (n = 10, N = 10, Supplementary Fig. 2).

### Fitting and statistical analysis

MATLAB was used to organize and normalize mG12 recruitment data, mean receptor expression data, mG12 expression data and prepare data for plotting in GraphPad Prism.

Python was used to perform linear fits in Supplementary Fig. 3, applying the least square method and an R^2^ > 0.98 criterion.

Final manuscript figures plotting, distribution fitting and statistics for data were carried out using GraphPad Prism Software (GraphPad, version 10.6.1). For statistical testing either an ordinary one-way ANOVA with Tukey’s post hoc test was performed for multisample comparisons, while an unpaired two-sided *t*-test was used for two-sample comparisons.

