## Supplementary Figures 1-16 for "Direct tensile force activates Adgrl3 in a tethered agonist-dependent manner"

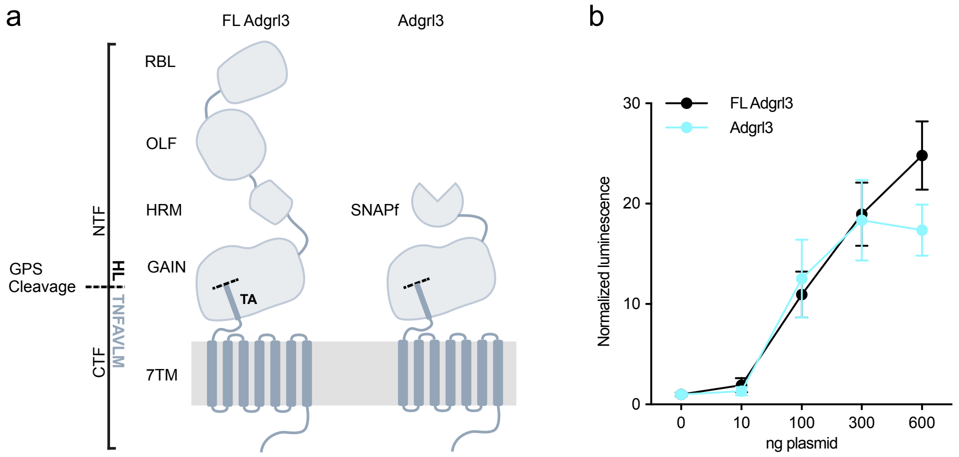


**Supplementary Fig. 1 | Structure and signaling activity of full length Adgrl3 and the construct used in this study.**

(**a**) Schematic of full-length (FL) Adgrl3 and the construct used in this study, in which the N-terminal region upstream of the GAIN domain is replaced by a SNAPf-tag (construct referred to as Adgrl3 throughout the manuscript). NTF, N-terminal fragment; CTF, C-terminal fragment; RBL, rhamnose-binding lectin; OLF, olfactomedin; HRM, hormone receptor motif; GPS, GPCR proteolytic site; GAIN, GPCR autoproteolysis-inducing domain; 7TM, seven-transmembrane domain. (**b**) FL Adgrl3 and Adgrl3 activate intracellular signaling at comparable levels, assessed using a luminescence-based serum response element (SRE) reporter assay to report on signaling activity downstream G protein activation. Data are normalized to empty vector control and shown as mean ± SEM. N = 3 independent experiments.

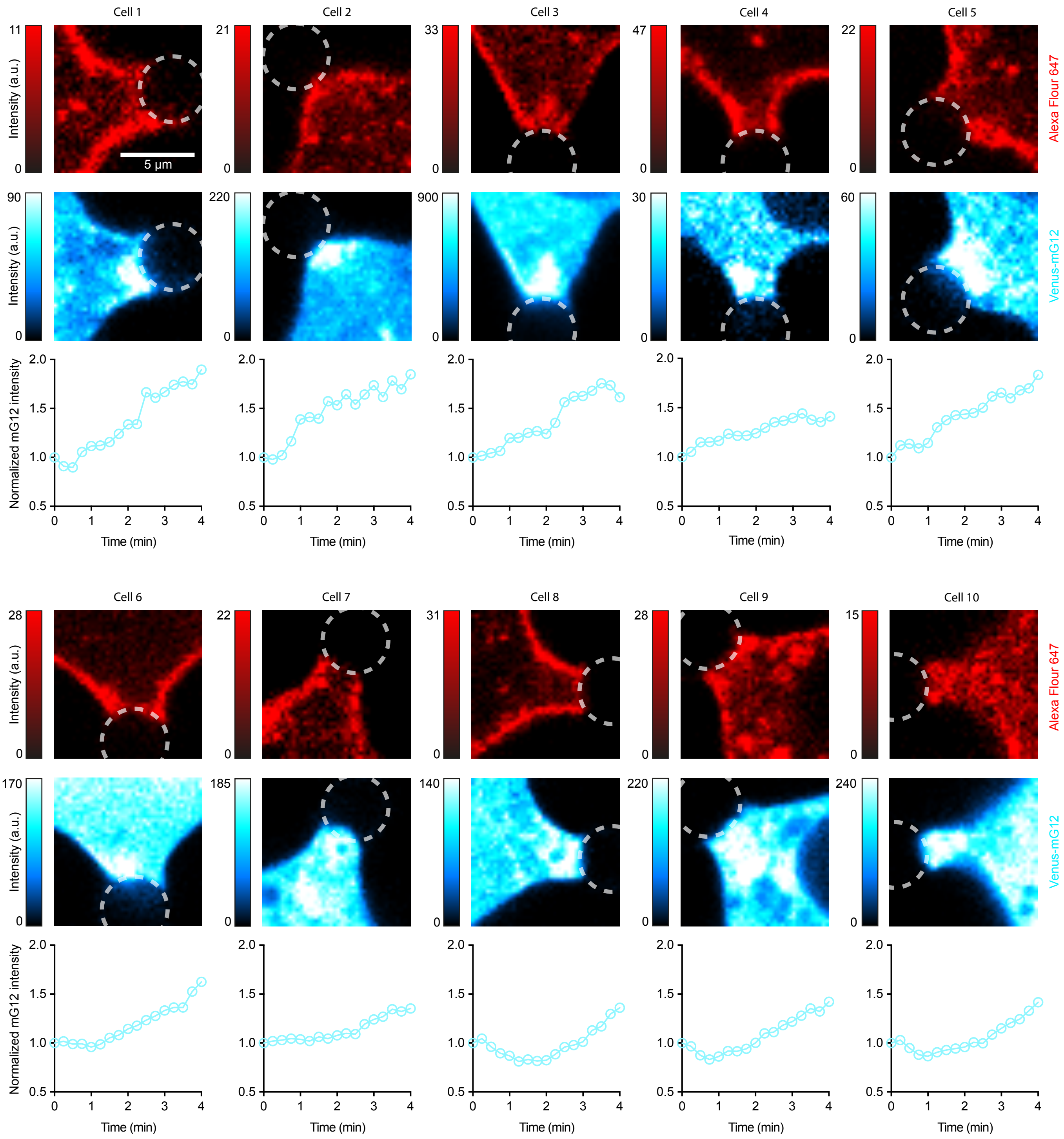


**Supplementary Fig. 2 | Individual cell deformations for Adgrl3- and mG12-expressing cells exposed to tensile forces.**
Confocal micrographs showing fluorescence from AF-647-labeled Adgrl3 (top row, red) and mG12 (bottom row, blue). Images show each of the 10 cells included in Fig. 1e at 4 min after onset of cell deformation. Circles (dotted) indicate bead position. Color scales represent intensity in arbitrary units and are adjusted individually for each micrograph. Scale bar, 5 µm. Changes in Venus-mG12 intensity are shown for each cell, with time zero defined as the onset of cell deformation. The methodology for quantifying Venus-mG12 intensities at the bead-cell interface is outlined in Supplementary Fig. 6. n = 10 cells from N = 10 independent experiments.

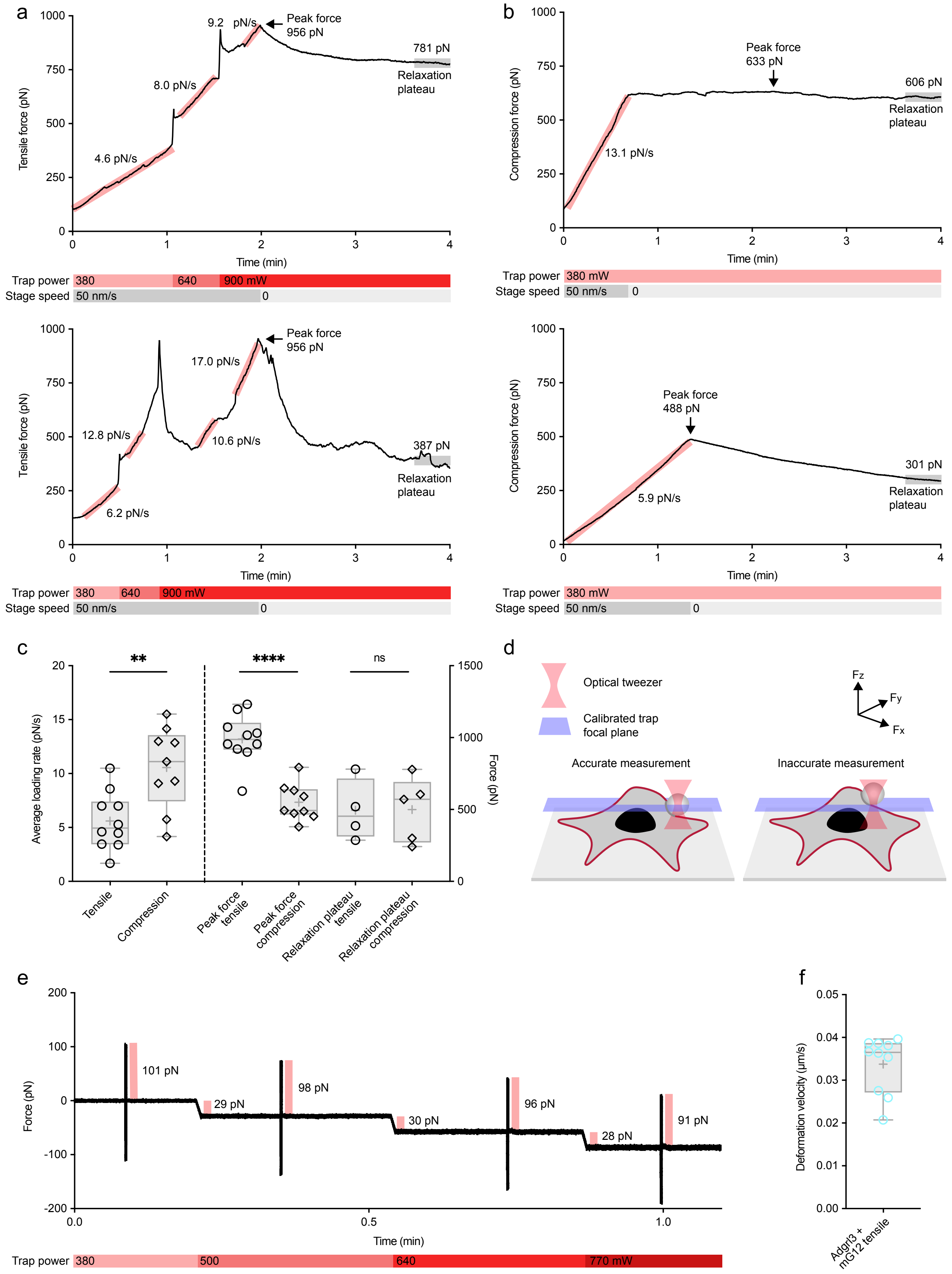


**Supplementary Fig. 3 | Quantification of tensile and compression forces.**
(**a-b**) Two exemplary force curves for application of tensile force (a) and compression force (b) to Adgrl3 over a 4-min time course from the onset of cell deformation (a) and onset of compression (b). Light red lines mark linear fits, and show the loading rates in different phases of the active force-loading region (large gray box), the black arrow marks the peak force and the gray bar the force plateau at 4 minutes. Under the exemplary force curves the laser power of the optical trap is shown together with the speed of the stage movement. For

the tensile force application force curves are largely linear but show two transient spikes from increased laser power required to maintain deformation at higher forces. In the first ~2 minutes of force application, we deform the cell at a velocity that exceeds the viscoelastic relaxation of the cell and produce increasing forces. In the last ~2 minutes of the force-application, we stopped stage motion, cells relax and we typically reach a plateau with constant force load. The measured average loading rates (c) are therefore distributed across the engaged receptors in the bead-cell contact area at any time. (**c**) Average force loading rate (left) and peak forces and plateaus (right) for both tensile and compression force application procedures. Circle: tensile; diamond: compression. n = 9-10 cells from N = 3 experiments for loading rate and peak force averages, and n = 4-5 cells from N = 3 experiments for plateau averages. Box plots indicate the median (central line), overall mean (central +), and interquartile range (lower and upper lines represent the 25th and 75th percentiles, respectively), while the whiskers are defined according to the Tukey criterion. (**d**) Schematic illustration showing potential force measurement inaccuracies caused by displacement of the bead along the z-axis from the calibrated focal plane of the optical trap during force application. Measurements where the bead showed such displacement were not included in the analysis. (**e**) Calibration experiment used to correct force offsets at each laser power (see Methods). The trap position oscillates in the x direction with an amplitude of 10 um and a frequency of 50 Hz for 0.2 seconds, for each tested laser power (15%, 20%, 25%, 30%). The force offset, measured as the average force during the first 3 seconds after the laser power reaches a new stable value, is subtracted from the corresponding force regime, to recover the correct force. Similar fluid drag forces due to the oscillation are reported for all laser powers tested, with up to 9% error. (**f**) Deformation velocities (see Methods) quantified for the 10 Adgrl3- and mG12-expressing cells exposed to tensile forces (Supplementary fig. 2). Open circles represent single cells, box plots indicate the median (central line), overall mean (central +), and interquartile range (lower and upper lines represent the 25th and 75th percentiles, respectively), while the whiskers are defined according to the Tukey criterion. Statistics: Unpaired two-tailed t-test between the pairs: **P = 0.0042, ****P < 0.0001, ns (not significant) P = 0.9692.

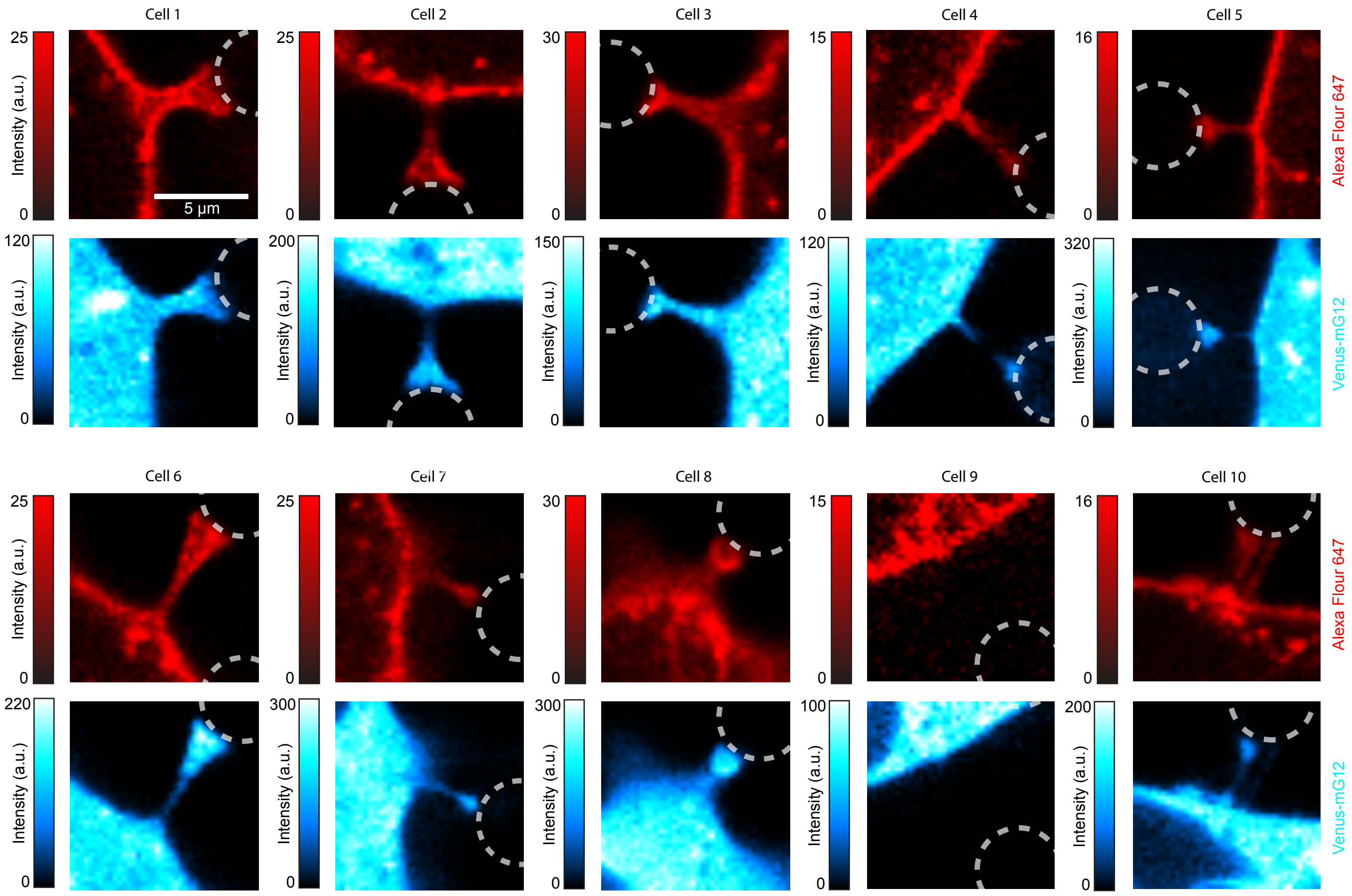


**Supplementary Fig. 4 | Cells exposed to tensile force application without BG-biotin result in thin tethers and no global deformations.**

Confocal micrographs showing AF-647-labeled Adgrl3 LM>AA (top row, red) and Venus-mG12 (bottom row, blue) in experiments performed without BG-biotin, preventing specific bead-receptor coupling. Images show 10 individual cells at 4 min following the same experimental protocol used for Fig. 1e and Supplementary Fig. 2. No large cell deformations were observed; only thin membrane tethers formed. Color scales represent intensity in arbitrary units. Scale bar, 5 µm. n = 10 cells from N = 4 independent experiments.

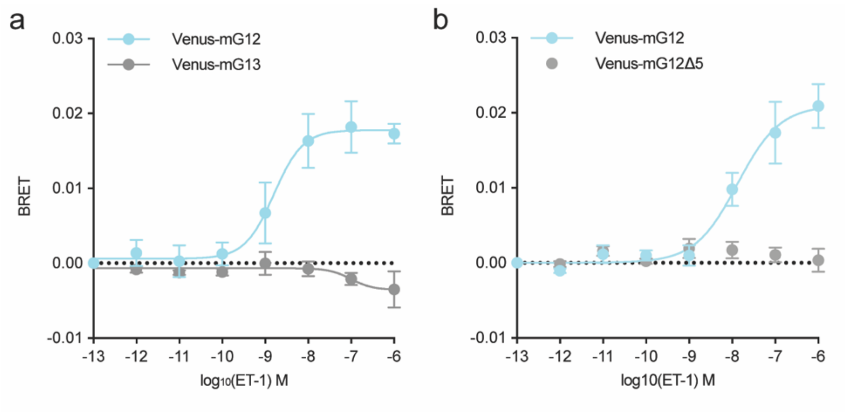


**Supplementary Fig. 5 | GPCR activation leads to recruitment of mG12, but not mG13.**
(**a**) Bioluminescence resonance energy transfer (BRET) ensemble assay measuring receptor activation-dependent recruitment of Venus-mG12 and Venus-mG13. BRET between the well-characterized Gq/G12/13-coupled receptor endothelin receptor type A (ETA), fused to a luminescent donor (Rluc8), and Venus-mG12 or Venus-mG13 (Acceptor) is plotted as a function of increasing concentrations of the ETA receptor agonist Endothelin 1 (ET-1). (**b**) BRET assay as in (a), showing recruitment of Venus-mG12 but not Venus-mG12Δ5. Data are normalized to buffer controls and show the BRET effect induced by ET-1, as mean ± SEM; N = 3 independent experiments.

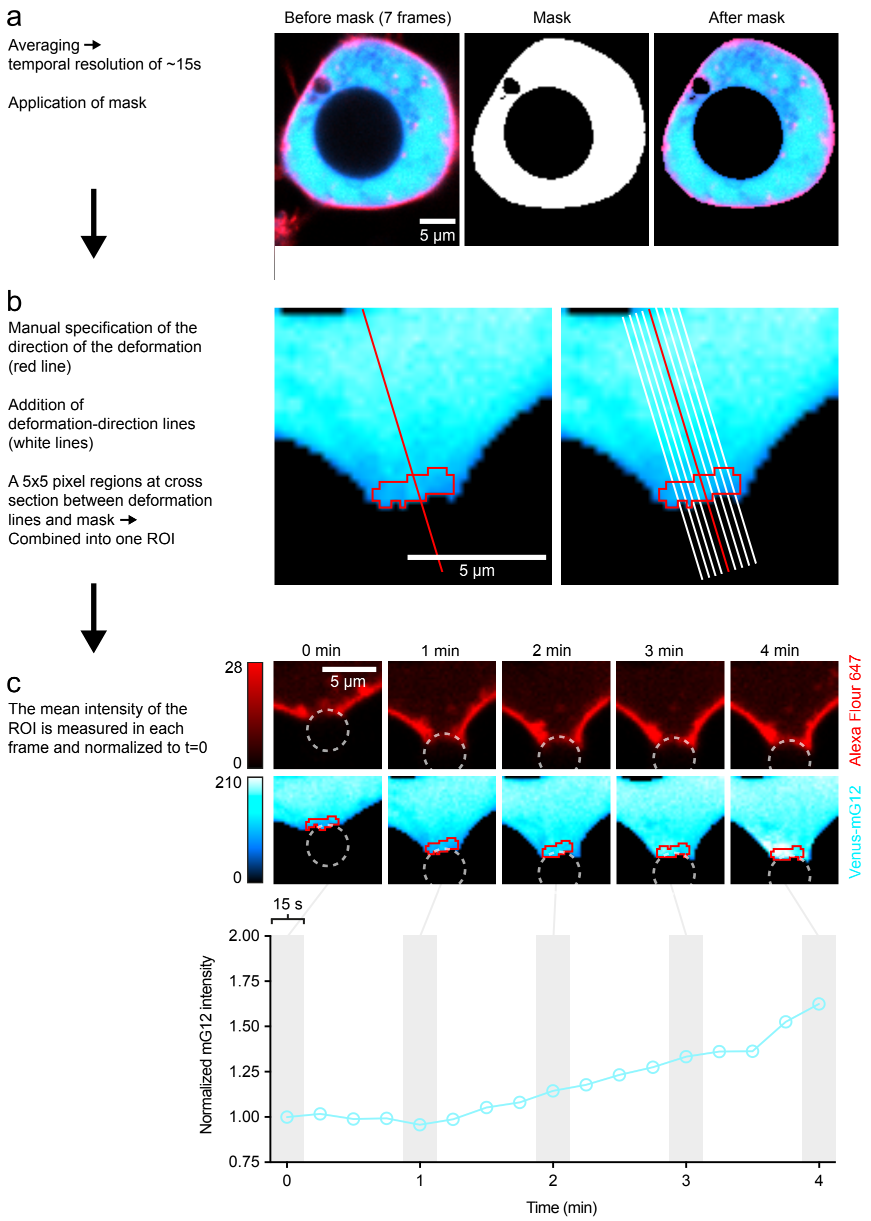


**Supplementary Fig. 6 | Quantification of mG12 recruitment.**
(**a-c**) The intensity quantification is performed using a custom-made macro for Fiji (Methods), and included the main steps shown in a-c. (a) To reduce noise, frames were averaged to a final temporal resolution of ~15s. A mask was then applied to define the location of the cell-bead interface and to reduce noise from background fluorescence. (b) The direction of the deformation was specified manually (red line). A number of parallel lines (white lines) were added with a distance of 1 pixel (250 nm), to span the entire width of the bead-cell interface, on both sides of the red line indicating the direction of deformation. For each time-averaged frame, a 5x5 pixel region was defined at the intersection between the deformation-direction lines and the mask defining the bead-cell interface shown in (a). All pixels in the 5x5 pixel regions were combined into one region of interest (ROI) spanning ~3-4 pixels (750-1000 nm) into the cell to include the entire intensity signal from the membrane-bound Venus-mG12 proteins and minimize contributions from non-specific effects from the optical trap (for example thermal effects and gradient forces) within intracellular regions. (c) For each frame the mean intensity of the ROIs was determined for the duration of the deforming experiment (4 minutes). t = 0 was defined as the beginning of the deformation and the ROI intensities were normalized to t = 0 for each cell.

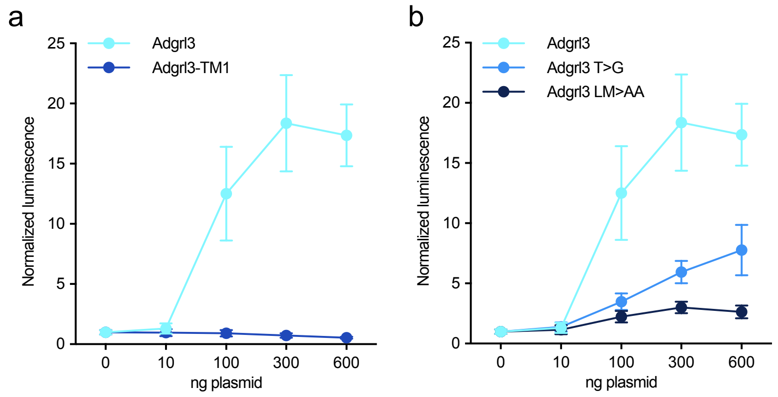


**Supplementary Fig. 7 | Signaling activity of Adgrl3 constructs.**
(**a**) SRE reporter assays showing that Adgrl3-TM1 does not elicit gene dose-dependent signaling, in contrast to Adgrl3. Constructs were expressed at comparable plasma membrane levels (see Supplementary Fig. 8). (**b**) SRE assays for Adgrl3, Adgrl3 T>G, and Adgrl3 LM>AA. TA mutations (LM>AA) strongly impair signaling, whereas the cleavage-deficient T>G mutant retains signaling activity. Adgrl3 and Adgrl3 LM>AA show comparable surface expression, while Adgrl3 T>G shows slightly reduced expression in ensemble measurements (Supplementary Fig. 8). Data are normalized to empty vector control and shown as mean ± SEM. N = 3 independent experiments.

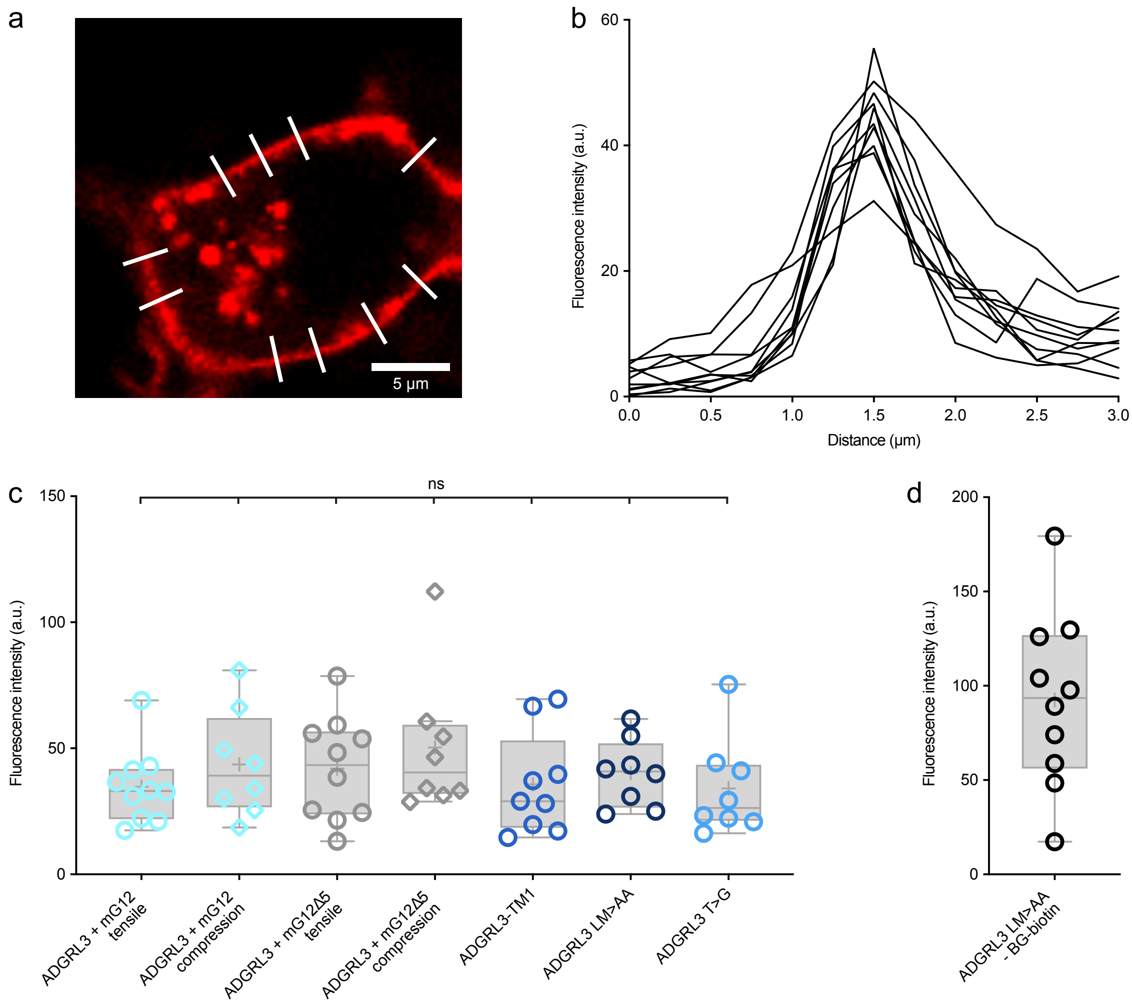


e

| Comparison | | Result | P value |
|---|---|---|---|
| Adgrl3 + mG12 tensile | Adgrl3 + mG12 compression | ns | 0,9652 |
| Adgrl3 + mG12 tensile | Adgrl3 + mG12Δ5 tensile | ns | 0,9841 |
| Adgrl3 + mG12 tensile | Adgrl3 + mG12Δ5 compression | ns | 0,6646 |
| Adgrl3 + mG12 tensile | Adgrl3-TM1 | ns | >0.9999 |
| Adgrl3 + mG12 tensile | Adgrl3 LM>AA | ns | 0,9974 |
| Adgrl3 + mG12 tensile | Adgrl3 T>G | ns | >0.9999 |
| Adgrl3 + mG12 compression | Adgrl3 + mG12Δ5 tensile | ns | >0.9999 |
| Adgrl3 + mG12 compression | Adgrl3 + mG12Δ5 compression | ns | 0,9942 |
| Adgrl3 + mG12 compression | Adgrl3-TM1 | ns | 0,9827 |
| Adgrl3 + mG12 compression | Adgrl3 LM>AA | ns | 0,9999 |
| Adgrl3 + mG12 compression | Adgrl3 T>G | ns | 0,9616 |
| Adgrl3 + mG12Δ5 tensile | Adgrl3 + mG12Δ5 compression | ns | 0,9747 |
| Adgrl3 + mG12Δ5 tensile | Adgrl3-TM1 | ns | 0,9936 |
| Adgrl3 + mG12Δ5 tensile | Adgrl3 LM>AA | ns | >0.9999 |
| Adgrl3 + mG12Δ5 tensile | Adgrl3 T>G | ns | 0,9813 |
| Adgrl3 + mG12Δ5 compression | Adgrl3-TM1 | ns | 0,7491 |
| Adgrl3 + mG12Δ5 compression | Adgrl3 LM>AA | ns | 0,9507 |
| Adgrl3 + mG12Δ5 compression | Adgrl3 T>G | ns | 0,6748 |
| Adgrl3-TM1 | Adgrl3 LM>AA | ns | 0,9993 |
| Adgrl3-TM1 | Adgrl3 T>G | ns | >0.9999 |
| Adgrl3 LM>AA | Adgrl3 T>G | ns | 0,9963 |

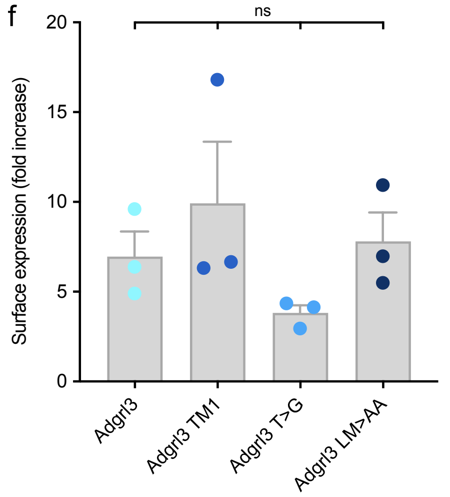


**Supplementary Fig. 8 | Adgrl3 constructs show identical surface expression levels across all optical tweezer experiments.**

(**a**) Example cell illustrating surface fluorescence quantification using AF-647-labeled Adgrl3. Line profiles were measured perpendicular to the cell membrane at ten different locations. Scale bar, 5 µm. (**b**) Intensity profiles from the line scans shown in (a). The mean of the maximum intensity from each of the ten line scans was used as the surface expression value for a given cell. (**c**) Surface expression levels for cells included in Fig. 1e and Fig. 2b and d (dots indicate individual cells). Box plots indicate the median (central line), overall mean (central +), and interquartile range (lower and upper lines represent the 25th and 75th percentiles, respectively), while the whiskers are defined according to the Tukey criterion. Ordinary one-way ANOVA (Tukey's multiple comparison test, with single pooled variance) is used to compare all conditions. P values shown in (e). (**d**) Surface expression for control cells from Supplementary Fig. 4 (no BG-biotin). (**e**) P values from the statistical comparisons in (c). (**f**) Ensemble surface expression measured using a plate reader (ClarioSTAR). AF-647 fluorescence from Tet-induced cells was normalized to non-induced controls. Data are shown as mean ± SD; N = 3 independent experiments. Statistics: One-way ANOVA with Tukey's multiple comparisons between groups at 4 min: not significant (ns) (Adgrl3 vs. Adgrl3 T>G, P = 0.7024,

Adgrl3 vs. Adgrl3 LM>AA, P = 0.9909, Adgrl3 vs. Adgrl3 TM1, P = 0.7388, Adgrl3 T>G vs. Adgrl3 LM>AA, P = 0.5418, Adgrl3 T>G vs. Adgrl3 TM1, P = 0.2248, Adgrl3 LM>AA vs. Adgrl3 TM1, P = 0.8787).

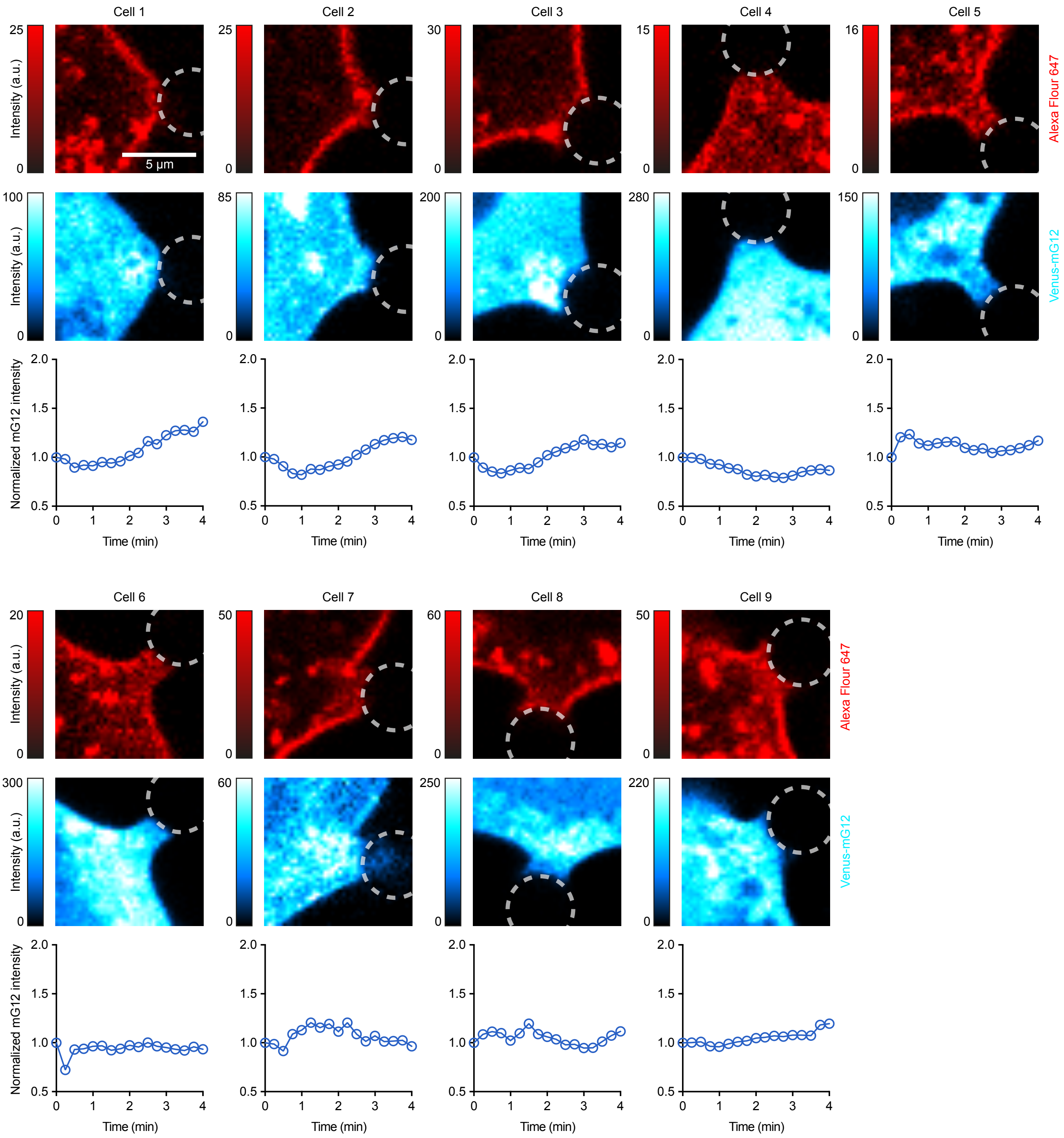


**Supplementary Fig. 9 | Individual cell deformations for Adgrl3-TM1 and mG12 expressing cells exposed to tensile forces.**
Confocal micrographs showing AF-647-labeled Adgrl3-TM1 (top row, red) and Venus-mG12 (bottom row, blue) for the cells included in Fig. 1e at 4 min after force application. Circles indicate bead position. Color scales represent intensity in arbitrary units. Scale bar, 5 µm. Changes in Venus-mG12 intensity are shown for each cell, with time zero defined as onset of cell deformation. The methodology for quantifying Venus-mG12 intensities at the bead-cell interface is outlined in Supplementary Fig. 6. n = 9 cells from N = 7 independent experiments. Intensity traces show that mG12 recruitment is not caused by endogenous receptors or by optical tweezer specific effects (i.e., thermal effects and gradient force).

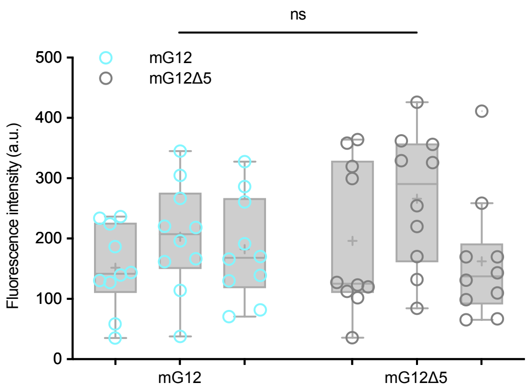


**Supplementary Fig. 10 | mG12 and mG12Δ5 are expressed at comparable levels.**
To quantify mG12 and mG12Δ5 expression levels, 10 cells expressing Adgrl3 LM>AA were imaged in three independent experiments. The intensity of Venus-mG12 was averaged across the whole cell and plotted as one datapoint. Box plots indicate the median (central line), overall mean (central +), and interquartile range (lower and upper lines represent the 25th and 75th percentiles, respectively), while the whiskers are defined according to the Tukey criterion. N = 3 individual experiments, n = 30 cells (10 for each experiment). No significant differences in expression levels were observed between mG12 and mG12Δ5. Statistics: Unpaired two-tailed t-test, ns (not significant) P = 0.3094.

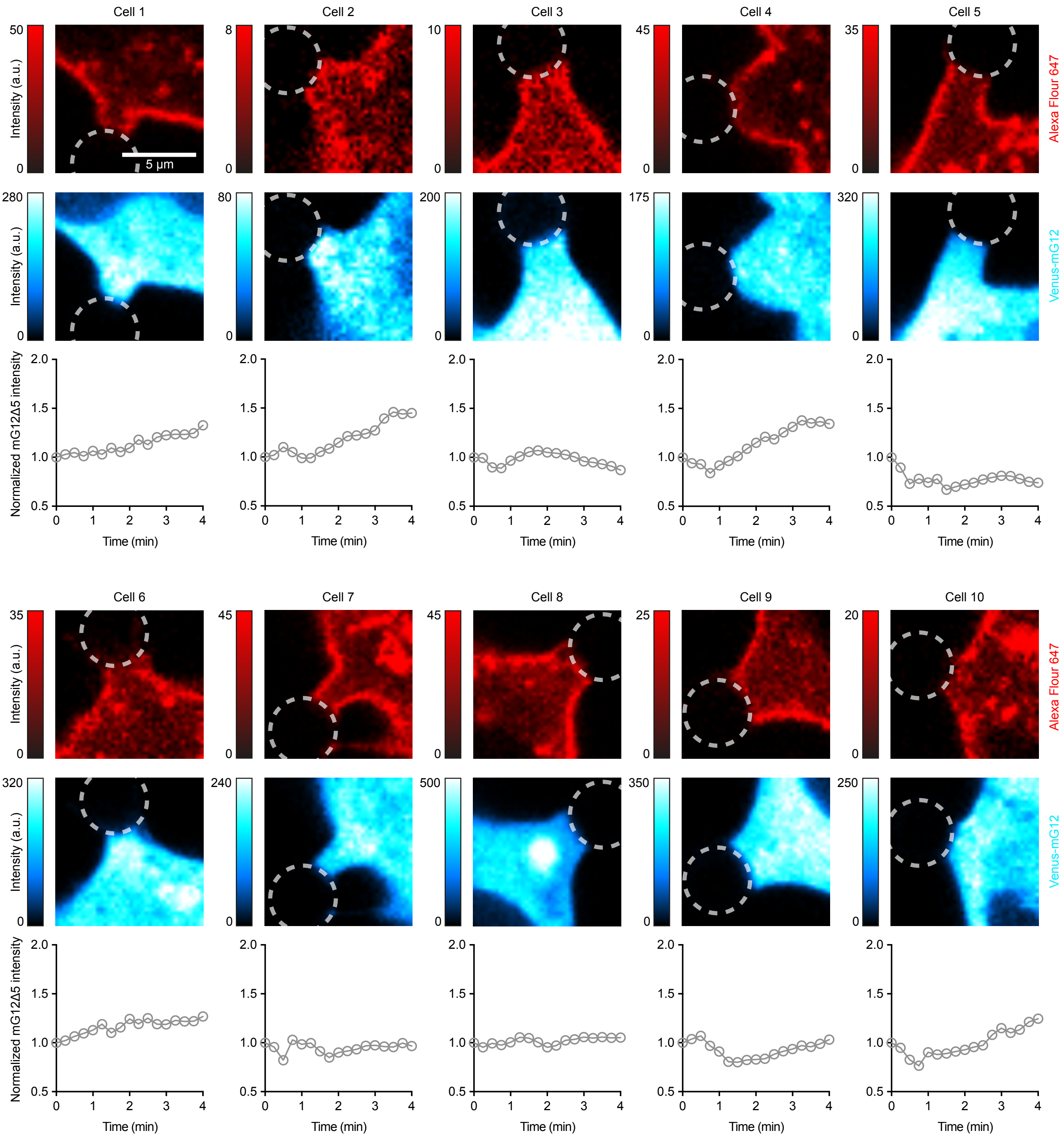


**Supplementary Fig. 11 | Individual cell deformation for Adgrl3 and mG12Δ5 expressing cells exposed to tensile forces.**
Confocal micrographs showing AF-647-labeled Adgrl3 (top row, red) and Venus-mG12Δ5 (bottom row, blue) for the cells included in Fig. 1e at 4 min after force application. Circles indicate bead position. Color scales represent intensity in arbitrary units. Scale bar, 5 µm. Changes in Venus-mG12 intensity are shown for each cell, with time zero defined as the onset of cell deformation. The methodology for quantifying Venus-mG12 intensities at the bead-cell interface is outlined in Supplementary Fig. 6. n = 10 cells from N = 10 independent experiments. Intensity traces show that mG12 recruitment is not caused by effects related to the optical tweezer (i.e., thermal effects and gradient force) or the change in global cellular geometry upon deformation. Some traces show a slight tendency in increased mG12 signal, but this effect is more global (see cell 4 and 10) and not at the edge of the bead-cell membrane (Supplementary Fig. 2), but rather in the vicinity of our defined ROI (Supplementary Fig. 6).

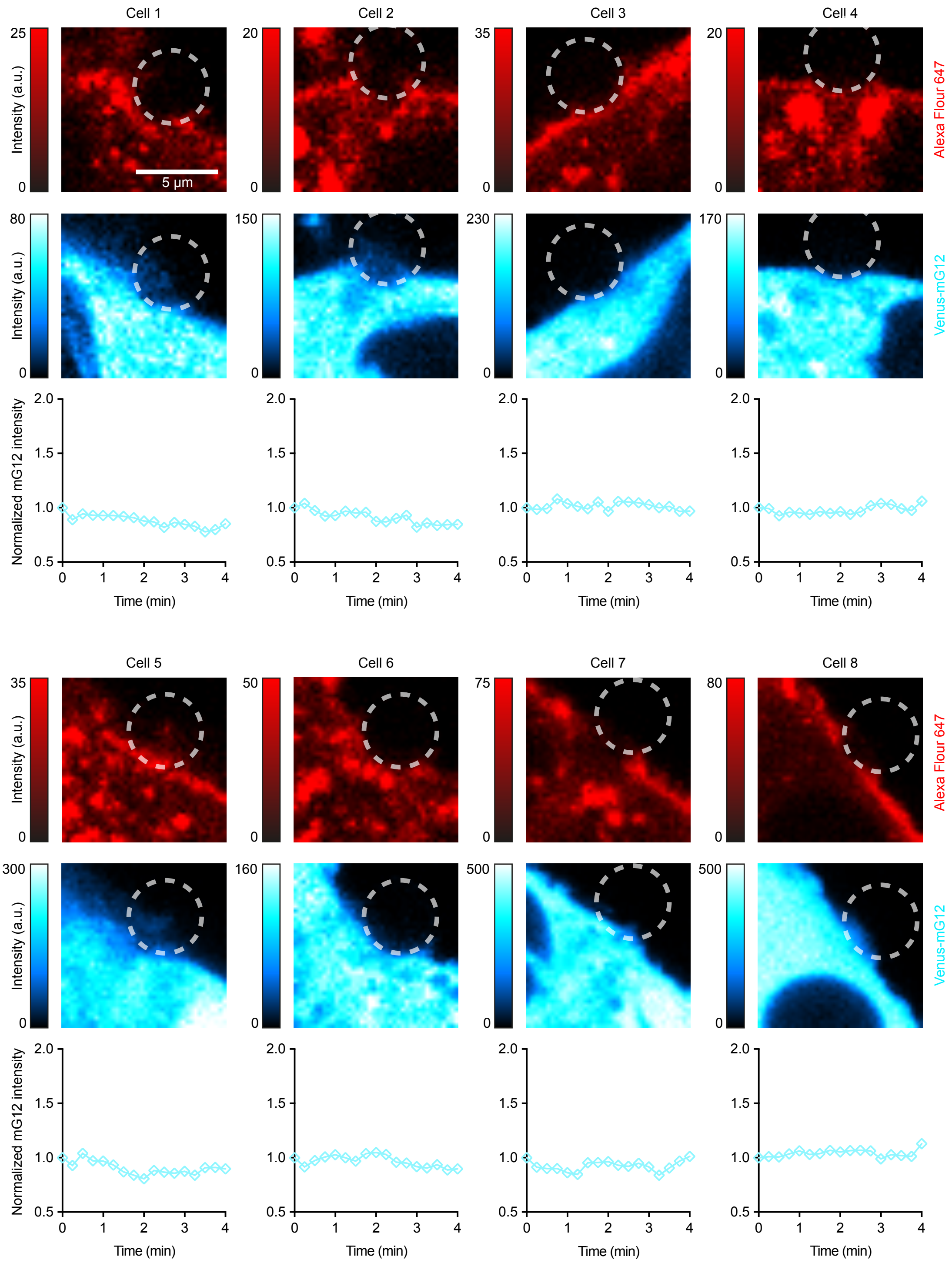


**Supplementary Fig. 12 | Individual cell deformations for Adgrl3 and mG12 expressing cells exposed to compression forces.**

Confocal micrographs from compression experiments showing AF-647-labeled Adgrl3 (top row, red) and Venus-mG12 (bottom row, blue) for the cells included in Fig. 2b at 4 min after compression onset. Circles indicate bead position. Color scales represent intensity in arbitrary units. Scale bar, 5 µm. Changes in Venus-mG12 intensity are shown for each cell, with time zero defined as the onset of cell compression. n = 8 cells from N = 6 independent experiments.

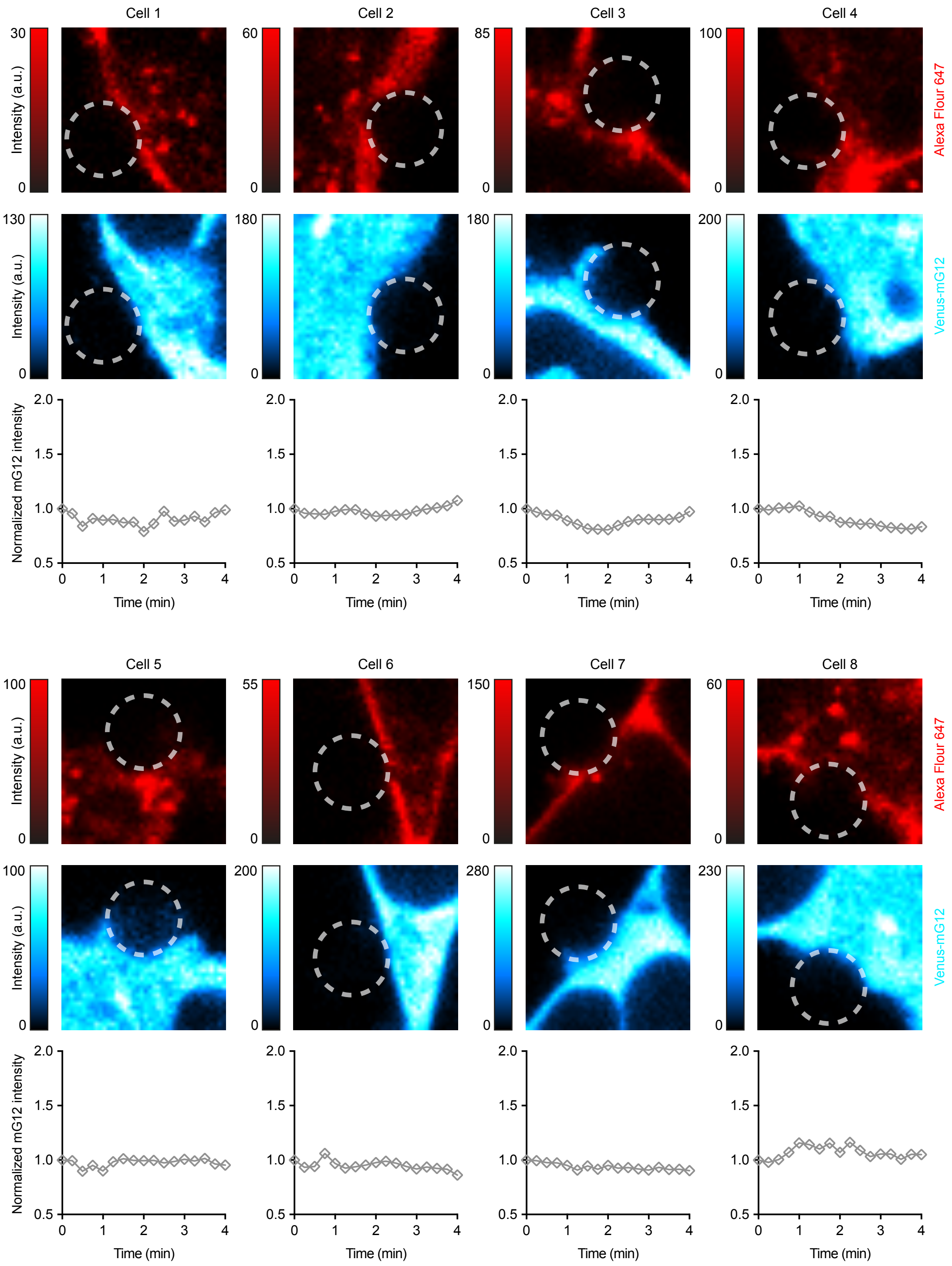


**Supplementary Fig | 13 Individual cell deformations for Adgrl3 and mG12Δ5 expressing cells exposed to compression forces.**

Confocal micrographs from compression experiments showing AF-647-labeled Adgrl3 (top row, red) and Venus-mG12Δ5 (bottom row, blue) for the cells included in Fig. 2 at 4 min after compression onset. Circles indicate bead position. Color scales represent intensity in arbitrary units. Scale bar, 5 µm. Changes in Venus-mG12Δ5 intensity are shown for each cell, with time zero defined as the onset of cell compression. n = 8 cells from N = 8 independent experiments.

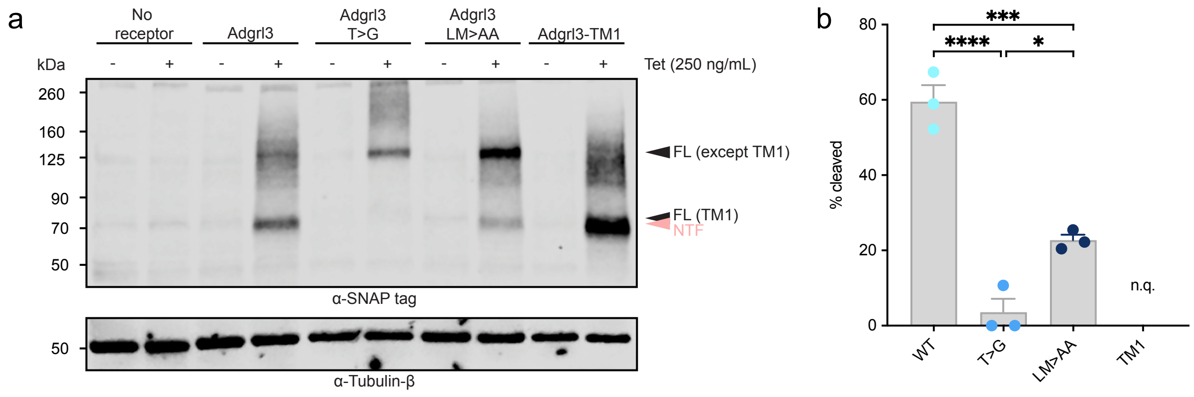


**Supplementary Fig | 14. Cleavage capacity of Adgrl3 constructs.**

(**a**) Western blot analysis of receptor expression following Tet induction (250 ng/mL), detected via the N-terminal SNAP tag. Bands corresponding to NTF and FL receptor are indicated; tubulin-β served as a loading control. (**b**) Quantification of receptor cleavage, expressed as the percentage of NTF relative to total receptor (NTF + FL). Cleavage of Adgrl3-TM1 could not be quantified (n.q.) due to insufficient resolution of NTF and FL bands. Data are shown as mean ± SEM with individual experiment values indicated. N = 3 independent experiments. Statistics: One-way ANOVA with Tukey's multiple comparisons: ***P = 0.0006, ****P < 0.0001, *P = 0.0164.

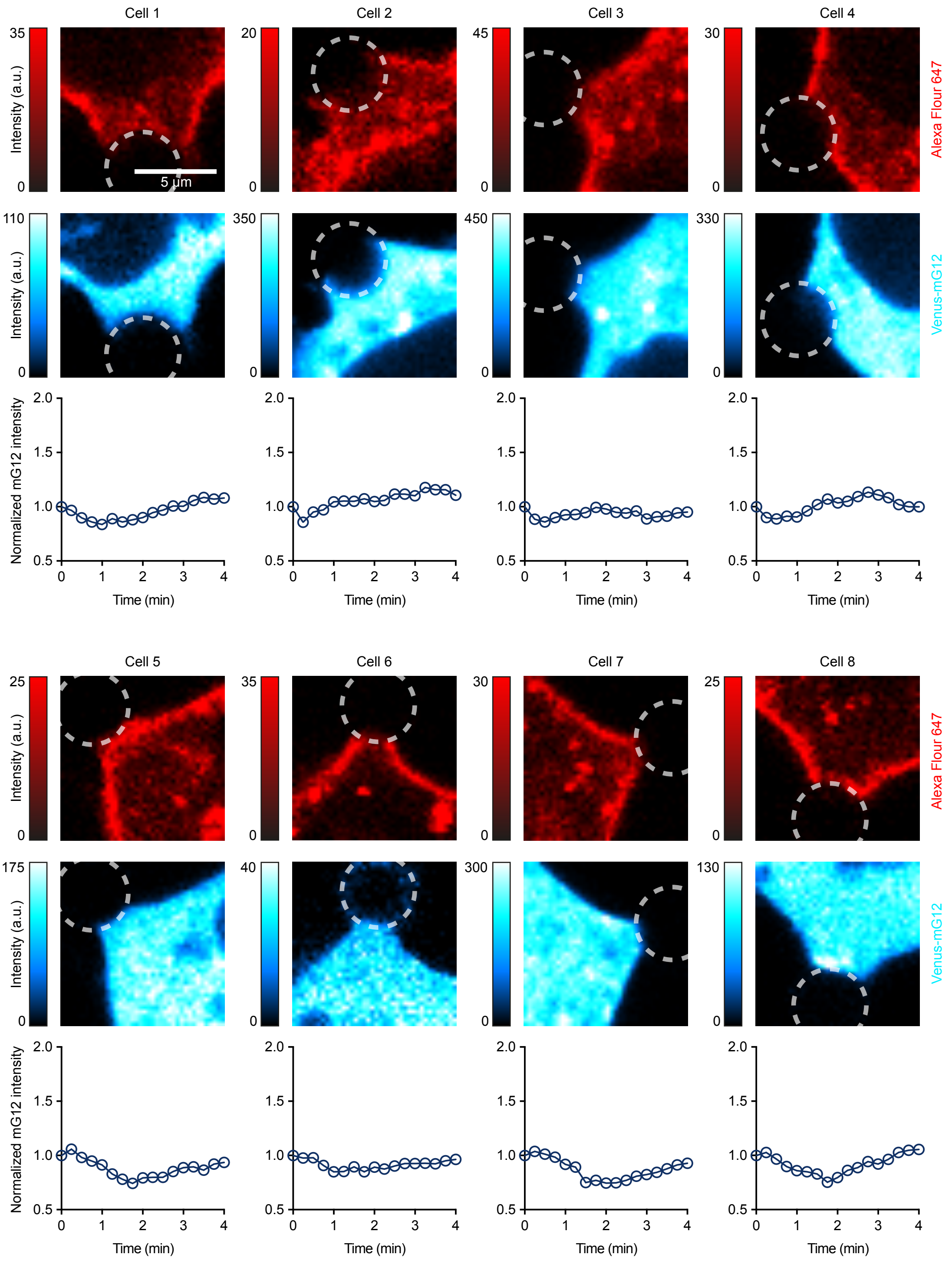


**Supplementary Fig. 15 | Individual cell deformations for Adgrl3 LM>AA and mG12 expressing cells exposed to tensile forces.**
Confocal micrographs showing AF-647-labeled Adgrl3 LM>AA (top row, red) and Venus-mG12 (bottom row, blue) for the cells included in Fig. 2d at 4 min after cell deformation onset. Circles indicate bead position. Color scales represent intensity in arbitrary units. Scale bar, 5 µm. Changes in Venus-mG12 intensity are shown for each cell, with time zero defined as the onset of cell deformation. n = 8 cells from N = 7 independent experiments.

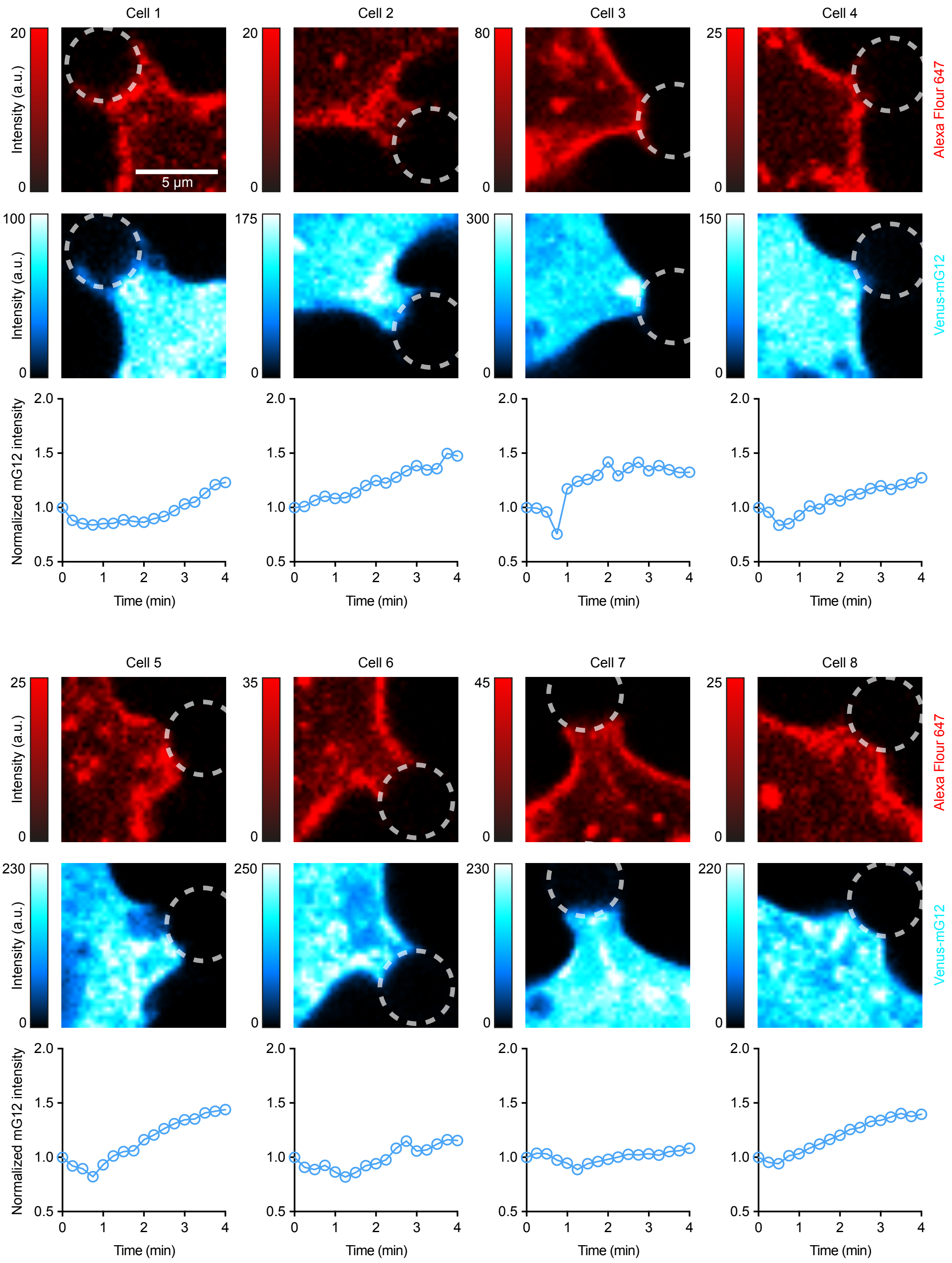


**Supplementary Fig. 16 | Individual cell deformations for Adgrl3 T>G and mG12 expressing cells exposed to tensile forces.**
Confocal micrographs showing AF-647-labeled Adgrl3 T>G (top row, red) and Venus-mG12 (bottom row, blue) for the cells included in Fig. 2d at 4 min after cell deformation onset. Circles indicate bead position. Color scales represent intensity in arbitrary units. Scale bar, 5 µm. Changes in Venus-mG12 intensity are shown for each cell, with time zero defined as the onset of cell deformation. n = 8 cells from N = 8 independent experiments.
